# NewroBus for the brain: humanized TfR1-targeting nanobodies with high BBB permeability and cargo transport capacity

**DOI:** 10.1101/2025.04.20.649139

**Authors:** Tao Yin, Metin Yesiltepe, Sanjay Metkar, Aubin Ramon, Matthew Greenig, Pietro Sormanni, Luciano D’Adamio

**Affiliations:** Department of Pharmacology, Physiology & Neuroscience New Jersey Medical School, Brain Health Institute, Jacqueline Krieger Klein Center in Alzheimer’s Disease and Neurodegeneration Research, Rutgers, The State University of New Jersey, 205 South Orange Ave, Newark, NJ, 0703, USA; Centre for Misfolding Diseases, Yusuf Hamied Department of Chemistry, University of Cambridge, Lensfield road, CB2 EW Cambridge, UK; NanoNewron LLC, Townsend Hall T27, 000 Morris Avenue, Union, NJ 07083

**Keywords:** TNFα, Nanobody, Blood-brain-barrier, Alzheimer Disease, TNFα inhibition, single-domain antibody

## Abstract

Effective delivery of therapeutics to the brain is restricted by the blood-brain barrier (BBB). A strategy to overcome this limitation involves taking advantage of receptor-mediated transcytosis pathways, such as those mediated by transferrin receptor 1 (TfR1), which is highly expressed on brain endothelial cells and naturally transports iron-bound transferrin across the BBB. To exploit this mechanism, we immunized camelids with human TfR1 and cloned 470 VHH nanobody sequences from their B cells. From this repertoire, 24 nanobodies (TfR1b-Nbs) were identified that bind human TfR1 on the cell membrane. These nanobodies were screened for binding to human TfR1, lack of interference with transferrin binding and TfR1-mediated iron uptake, and the ability to cross the BBB via human TfR1-mediated transcytosis in newly generated humanized *Tfr1^h^* knock-in rats.

To improve developability and reduce potential immunogenicity, selected TfR1b-Nbs were humanized and optimized with computational and artificial intelligence (AI) algorithms, enhancing humanness, solubility, and VHH-nativeness. Eight optimized TfR1b-Nbs retained BBB permeability and were fused to humanized anti-TNFα nanobody inhibitors (TNFI-α or TNFI-β), generating 16 heterodimers. Fusion to these TNFIs served as a functional readout, confirming that TfR1b-Nbs can shuttle biologically active, BBB-impermeable payloads into the central nervous system (CNS). All heterodimers demonstrated CNS delivery after intravenous administration, and selected constructs also reached the brain via subcutaneous injection, maintaining high serum and cerebrospinal fluid (CSF) levels for up to 72 hours. A pilot study with one heterodimer showed that chronic administration in rats humanized for both transferrin and TfR1 caused no hematological toxicity or signs of anemia – a key safety concern when targeting TfR1. These results establish humanized TfR1b-Nbs – designated *NewroBus* – as promising BBB shuttles for the safe and effective therapeutic delivery of biologics to the brain.

## Introduction

Targeting TfR1 to enhance drug delivery across the BBB has emerged as a rapidly advancing area of therapeutic development. This strategy leverages the fact that TfR1 is highly expressed on brain endothelial cells and its transcytosis activity can be exploited to deliver therapeutic molecules into the CNS ^1–3^. Several companies are actively pursuing distinct TfR1-mediated strategies to enable CNS penetration of biologics. Notable examples include Denali Therapeutics, which developed the Transport Vehicle (TV) platform utilizing engineered Fc domains that bind TfR1 to drive receptor-mediated transcytosis ^4^; Roche, with its Brain shuttle technology, based on a bispecific antibody format ^5^; and Apertura Gene Therapy, which employs engineered AAV capsids specifically designed to target human TfR1 and improve CNS gene delivery ^6^. BioArctic has also introduced its Brain Transporter™ technology, although mechanistic and efficacy data remain unpublished.

Camelids naturally produce heavy-chain-only antibodies that lack light chains and consist of a single variable domain (VHH) followed by constant domains CH2 and CH3 ^7^. The isolated VHH domains, commonly referred to as nanobodies (Nbs), are compact, stable antigen-binding fragments with a typical molecular weight of 12–14 kDa. Compared to conventional monoclonal antibodies (mAbs), nanobodies offer several advantages: (1) their convex paratopes allow binding to recessed or cryptic epitopes, such as receptor binding pockets that are often inaccessible to the flat paratopes of mAbs; (2) they exhibit high stability across a wide pH range; (3) they have low immunogenicity, which can be further minimized through humanization or deimmunization; and (4) their small size and structural robustness make them more amenable to alternative delivery routes, including potentially oral administration. Importantly, nanobodies targeting transferrin receptor 1 (TfR1) have been shown to mediate transcytosis across the BBB ^8,9^, highlighting their potential as molecular buses. Additionally, nanobody-based TfR1 binders are monovalent and thus less likely to interfere with the physiological function of TfR1 in iron transport. Unlike bivalent antibodies, they are less likely to disrupt the transferrin (TF)–TfR1 interaction essential for cellular iron uptake or to promote TfR1 endocytosis, which typically requires engagement of both subunits of the TfR1 dimer. This minimizes the risk of iron deficiency-related toxicities, an important consideration in the development of TfR1-targeted therapeutics.

Based on these properties, we set out to identify and characterize novel anti-TfR1 nanobodies capable of efficiently crossing the BBB, with the long-term goal of using them as “Trojan horses” to deliver otherwise BBB-impermeable therapeutics into the brain.

## Results

### Identification of TfR1b-Nbs

470 individual VHH domain clones isolated from PBMCs from 1 llama and 1 alpaca immunized with human TfR1 extracellular domain (Sino Biological, HPLC-11020-H07H) were tested for antigen binding. 106 unique α-TfR1-Nb VHH sequences were identified. The above experiments were performed at Abcore.

HEK293 human cells were transfected with a vector co-expressing human transferrin receptor 1 (hTfR1) and EGFP. Of the 106 α-TfR1 nanobodies (Nbs) produced in bacteria, 24 bind to cell-surface hTfR1 on transfected cells (**Supporting Figures S1**, **S2**, **S3** and **S4**). These nanobodies were designated TfR1b-Nbs (TfR1-binding nanobodies). None of the 24 TfR1b-Nbs cross-reacted with rodent (rat or mouse) Tfr1 (**Supporting Figures S5**, **S6**, **S7** and **S8**). Based on the complementarity-determining regions (CDRs) identities, these 24 TfR1b-Nbs were grouped into ten families (**Figure 1A**).

**Figure 1.**
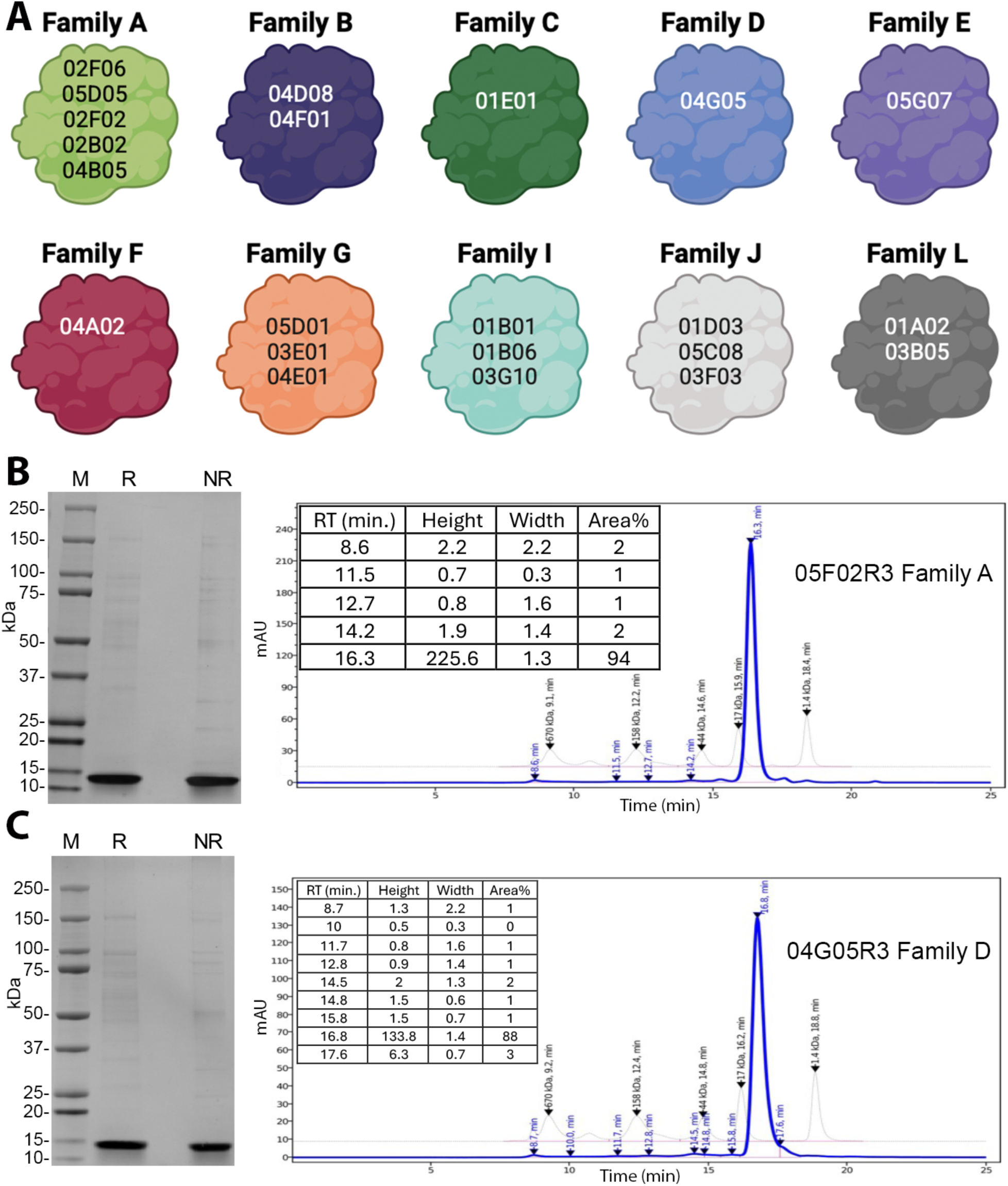
Protein purification and analysis of selected TfR1b-Nbs. **A)** Schematic representation of the 24 TfR1b-Nbs produced in mammalian cells. **B)** SDS-PAGE and SEC-HPLC analysis of TfR1b-Nb 02B02R3 (Family A). Coomassie-stained SDS-PAGE shows M = molecular weight marker; Lane 1 = purified 02B02R3 under reducing conditions; Lane 2 = under non-reducing conditions. Left panel: SEC-HPLC profile of purified 02B02R3. **C)** SDS-PAGE and SEC-HPLC analysis of TfR1b-Nb 04G05R3 (Family D), shown as in (B).

### Assessing the binding of TfR1b-Nbs produced in mammalian cells to human TfR1

Chinese Hamster Ovary (CHO) cells are a preferred system to produce therapeutic proteins due to their ability to generate complex, properly folded proteins with significantly lower endotoxin levels compared to bacterial expression systems, enhancing both the safety and clinical suitability of the final product. CHO-derived biologics have a well-established track record of regulatory approval, making this platform highly reliable for biopharmaceutical manufacturing.

The data in **Table 1** and **Figure 1B** and **1C** (showing representative results for 05F02R3 from Family A and 04G05R3 from Family D) demonstrate the high purity and yield of TfR1b-Nbs produced with transient transfection in CHO-S cells, supporting their suitability as a production platform for preclinical and clinical development. Protein production was performed at GenScript.

**Table 1.** Production of TfR1b-Nbs in mammalian cells was performed as for TNFI-Nbs. The table 1 shows quantity and purity of TfR1b-Nb purified from 100 ml culture supernatants.

| Name | Family | Conc. (mg/ml) | Purity SDS-PAGE (%) | Purity SEC-HPLC (%) | Endotoxin Level (EU/mg) | Total (mg) |
| --- | --- | --- | --- | --- | --- | --- |
| 02F06R3 | A | 0.45 | 75 | 92 | 0.111 | ~6.525 |
| 05D05R3 | A | 0.44 | 85 | 94 | 0.114 | ~6.38 |
| 05F02R3 | A | 0.46 | 90 | 94 | 0.109 | ~6.67 |
| 02B02R3 | A | 0.52 | 90 | 95 | 0.096 | ~7.54 |
| 04B05R3 | A | 0.53 | 50 | 96 | 0.094 | ~7.685 |
| 04D08R3 | B | 0.46 | 95 | 91 | 0.109 | ~6.67 |
| 04F01R3 | B | 0.42 | 80 | 89 | 0.119 | ~6.09 |
| 04G05R3 | D | 0.42 | 90 | 88 | 0.119 | ~6.09 |
| 05G07R3 | E | 0.29 | 70 | 83 | 0.172 | ~2.75 |
| 04A02R3 | F | 0.67 | 70 | 88 | 0.075 | ~9.045 |
| 05D01R3 | G | 0.6 | 75 | 89 | 0.083 | ~8.4 |
| 03E01R3 | G | 0.81 | 80 | 90 | 0.062 | ~11.34 |
| 04E01R3 | G | 0.88 | 70 | 92 | 0.057 | ~12.58 |
| 05D04R3 | J | 0.36 | 85 | 95 | 0.139 | ~5.04 |
| 01D03R3 | J | 0.84 | 75 | 94 | 0.064 | ~12.18 |
| 05C08R3 | J | 0.68 | 70 | 91 | 0.074 | ~9.18 |
| 01B03R3 | J | 0.73 | 80 | 94 | 0.068 | ~10.22 |
| 03F03R3 | J | 0.79 | 90 | 93 | 0.063 | ~11.06 |
| 01B01R3 | I | 0.92 | 95 | 96 | 0.054 | ~12.88 |
| 01B06R3 | I | 0.58 | 75 | 86 | 0.086 | ~8.7 |
| 03G10R3 | I | 0.79 | 85 | 88 | 0.063 | ~11.85 |
| 03B05R3 | L | 0.85 | 85 | 84 | 0.059 | ~12.75 |
| 01A02R3 | L | 0.72 | 70 | 83 | 0.069 | ~10.8 |

The biological activity of CHO-S–produced TfR1b-Nbs was assessed by testing their binding to human TfR1. HEK293 cells transfected with hTfR1 and EGFP showed strong staining with TfR1b-Nbs from Families A, B, D, G, and I, but not from Families E, F, J, or L (**Figure 2**). Similar results were obtained using CHEK-ATP089, a stable HEK293/hTfR1 cell line, where TfR1 expression was confirmed by transferrin-FITC staining (**Figure 3**).

**Figure 2.**
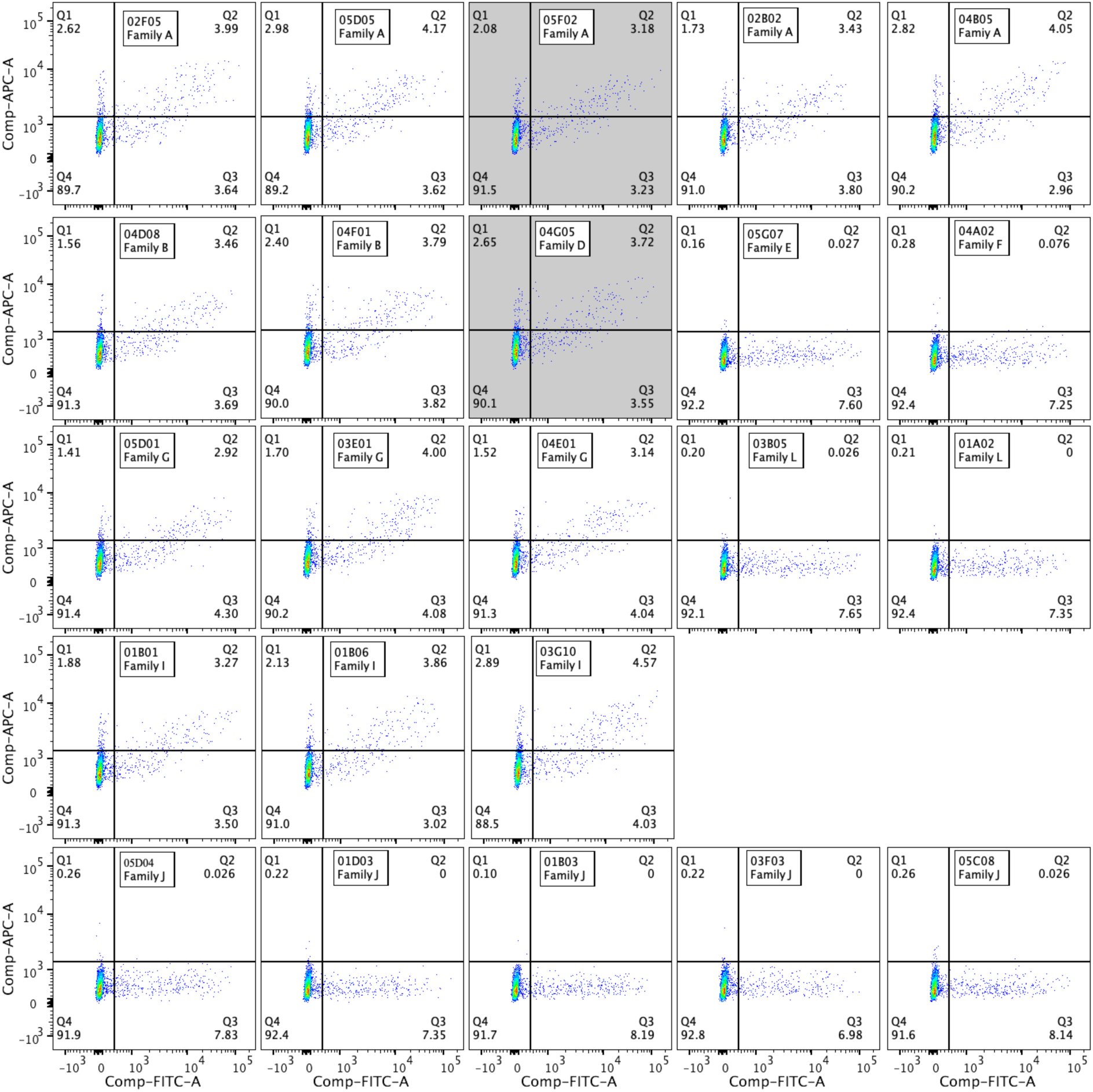
TfR1b-Nbs produced by mammalian cells bind to human TfR1. HEK293 cells were transfected with a vector expressing human TfR1 (alongside EGFP). Subsequently, cells were treated with each TfR1b-Nb at a concentration of 400 nanomolar, followed by incubation with an anti-His-APC antibody. The two nanobodies highlighted in grey are the parental sequences from which the final selected NewroBus molecules were derived.

**Figure 3.**
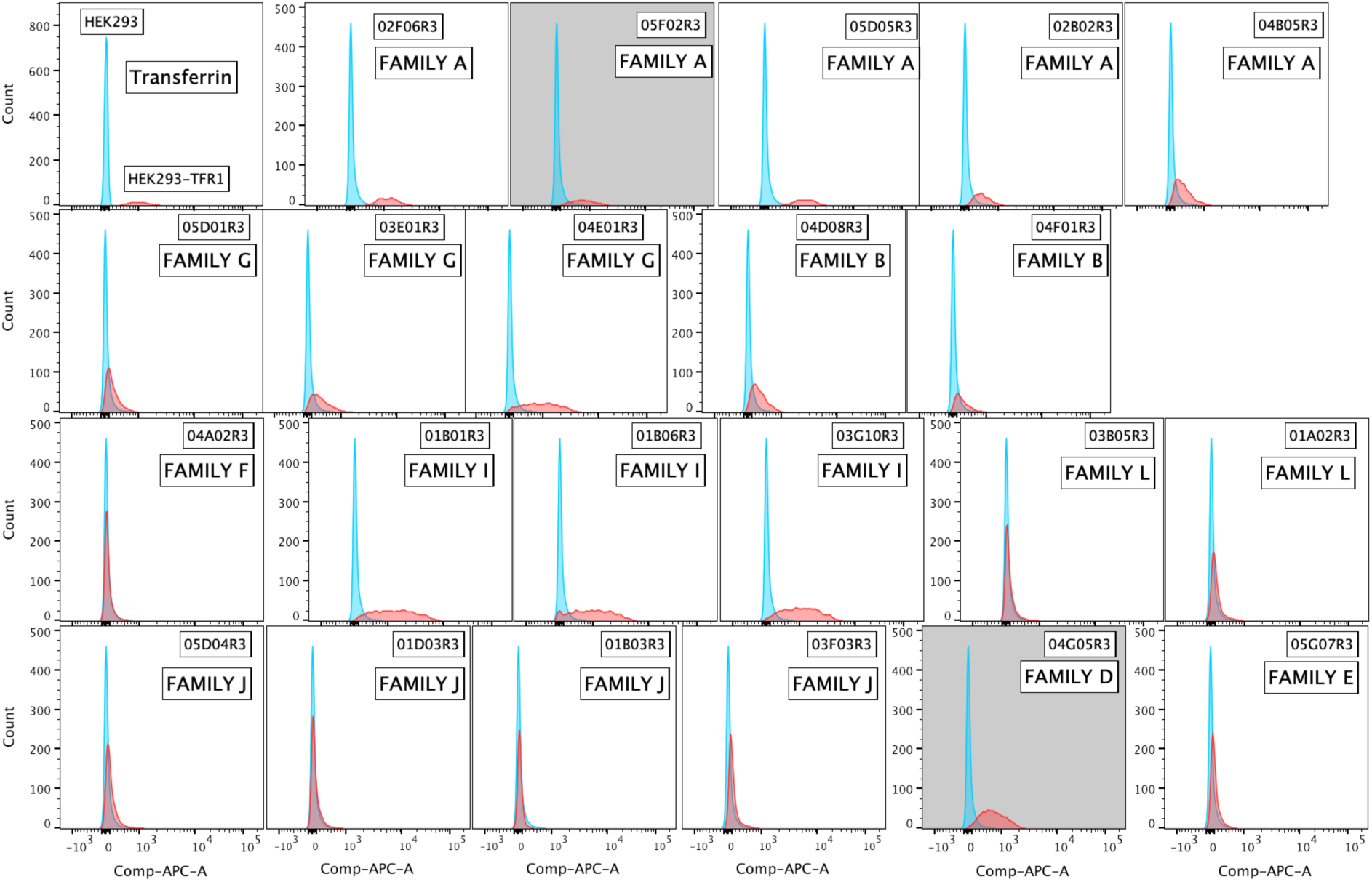
TfR1b-Nbs produced by mammalian cells bind to human TfR1. The upper-left panel shows the specific binding of human Transferrin-FITC (at a concentration of 2.5 nanomolar) to CHEK-ATP089 cells, with no binding observed on HEK293 cells. The remaining panels depict the binding of each TfR1b-Nb at a concentration of 40 nanomolar to CHEK-ATP089 cells. No binding was observed on HEK293 cells. The two nanobodies highlighted in grey are the parental sequences from which the final selected NewroBus molecules were derived.

### Identifying TfR1-Nbs compatible with transferrin-mediated iron transport

TF is the major ferric iron transport protein, which binds ferric (Fe3^+^) ions. The iron binding affinity of transferrin is pH dependent. In a neutral pH environment, TF (apotransferrin) binds iron with high affinity to form iron-bound TF (holotransferrin). In an acidic pH environment, the affinity of iron bound to transferrin decreases, dissociating iron from holotransferrin and releasing it into the environment. The importance of holotransferrin is to sequester Fe3+ ions in a relatively nonreactive and inert state to ensure normal free iron homeostasis in the body. Holotransferrin delivers iron to cells by binding to transferrin receptor (TfR1 or TfR2). Neutral pH at the cell surface promotes binding of holotransferrin to TfR1/TfR2. The receptor-ligand complex enters the cell through receptor-mediated endocytosis and is internalized into endosomes. Relatively lower endosomal pH results in the release of iron. The receptor-ligand complex is recycled to the cell surface, where apotransferrin dissociates from TfR1/TfR2 ^10^. Interference with these processes might lead to unintended toxic effects in patients treated with therapeutics based on TfR1b-Nbs.

To evaluate whether TfR1b-Nbs interfere with TF binding or uptake, we used CHEK-ATP089 cells. As expected, unlabeled TF competes with TF-FITC for receptor binding in a dose-dependent manner—effective at TF/TF-FITC ratios of 3 and 1, but not 0.3—confirming assay sensitivity (**Figure 4**). Using the same setup, we tested whether TfR1b-Nbs (2.5 µM) could reduce TF-FITC binding at a 1:1 molar ratio. All three Family I TfR1b-Nbs reduced TF-FITC binding, while none of the other 11 tested Nbs (from Families A, B, D, and G) showed interference (**Figure 4**).

**Figure 4.**
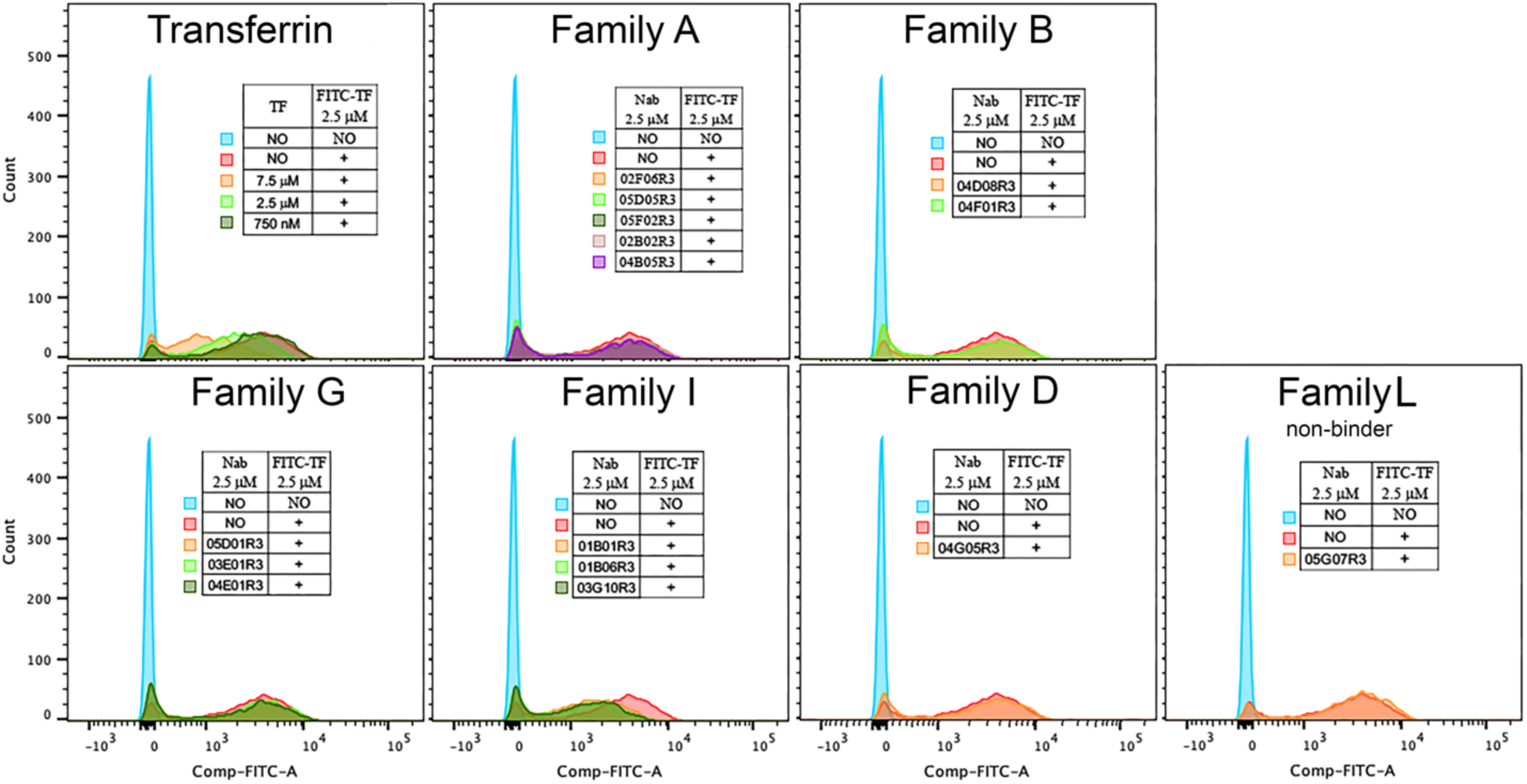
Selection of TfR1b-Nbs that do not interfere with TF-TfR1 interaction. The upper-left panel demonstrates the dose-dependent competition of unlabeled Transferrin for binding to TfR1 on the cell surface of CHEK-ATP089 cells in the presence of FITC-Transferrin. Subsequent panels assess the inhibitory activity of 14 TfR1b-Nbs from Families A, B, D, G, and I. A nanobody from Family L (which lacks binding to TfR1 on CHEK-ATP089 cells) serves as a negative control. In these experiments, CHEK-ATP089 cells were incubated with the specified proteins for 1 hour on ice before FACS analysis. Human Transferrin-FITC and TfR1b-Nbs were utilized at a concentration of 2.5 micromolar.

We next assessed whether TfR1b-Nbs affect TF uptake, using pHrodo Red-TF and live-cell imaging on the Incucyte system. pHrodo Red fluorescence increases in acidic compartments, providing a readout of endocytic uptake. Unlabelled TF inhibited pHrodo Red-TF uptake in a dose-dependent manner, even at high pHrodo Red-TF/TF ratios (312.5 nM vs. 30 nM) (**Figure 5**). Despite potential imprecision in calculating pHrodo Red-TF molarity, these results confirm that competition can impair uptake. Consistent with binding data, Family I Nb 01B01R3 significantly reduced pHrodo Red-TF uptake (**Figure 5**), though less potently than 30 nM TF. TfR1b-Nbs from Families A, B, D, and G had no impact (**Figure 5**). A Family J Nb, which does not bind TfR1, also showed no effect (**Figure 5**).

**Figure 5.**
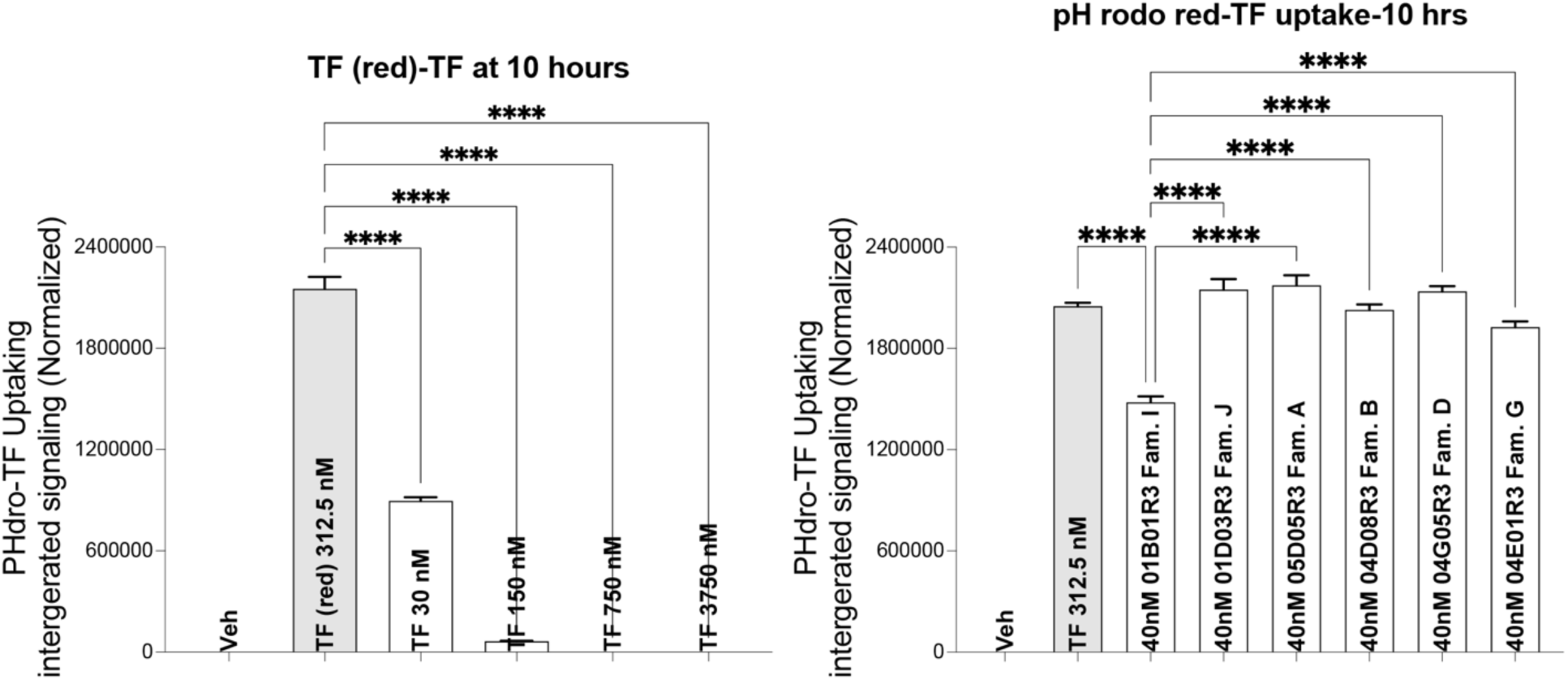
Selection of TfR1b-Nbs that do not interfere with TF uptake. The panel demonstrates the dose-dependent competition of unlabeled Transferrin pHrodo Red-TF uptake by CHEK-ATP089 cells. The right panel assess the inhibitory activity of representative TfR1b-Nbs from Families A, B, D, G, and I. A nanobody from Family J (which lacks binding to TfR1 on CHEK-ATP089 cells) serves as a negative control.

Based on these results, TfR1b-Nbs from Families A, B, D, and G appear most suitable for further development, as they do not interfere with TF-TfR1 interactions and iron uptake.

### Assessing in vivo BBB permeability of TfR1b-Nbs

Because TfR1b-Nbs do not bind rodent TfR1 (**Supporting Figures S5-S8**), rodents are unsuitable for in vivo BBB permeability studies. To overcome this problem, we generated knock-in (KI) rats in which the endogenous rat *Tfr1* gene was replaced with the human *TfR1* gene (*Tfr1^h^*). The generation and characterization of these KI rats will be described in a separate publication. These *Tfr1^h^* rats are viable, fertile, and exhibit normal reproductive behavior. Interbreeding of heterozygous *Tfr1^h^*animals produces *Tfr1^h/w^, Tfr1^w/w^* and *Tfr1^h/h^* offspring in expected Mendelian ratios, indicating that the humanized *Tfr1* allele does not impair embryonic development or viability. Furthermore, both male and female *Tfr1^h/h^* rats are fertile and can produce litters when crossed, confirming preserved reproductive capacity.

To assess whether humanization of *TfR1* compromises BBB integrity, we analyzed protein distribution between serum and CSF using SDS-PAGE in rats carrying humanized *TfR1* and *Tf* alleles. This analysis revealed no major differences in CSF protein content or pattern across genotypes. Rats with heterozygous or homozygous humanized *TfR1*(*Tfr1^h/w^* and *Tfr1^h/h^*), as well as double-humanized *Tf*/*TfR1* rats (*Tf^h/h^*:*Tfr1^h/h^*), exhibited CSF profiles similar to wild-type controls, indicating that BBB integrity is preserved following humanization (**Figure 6A**). A ∼15 kDa band of unknown identity was observed in the CSF of one control and one double-humanized rat. Serum samples (diluted 1:10) showed comparable protein patterns across all groups. Additional analysis of *Tfr1^h/h^* rats confirmed these findings, with only a faint ∼70 kDa band detectable in CSF, consistent with intact BBB function and normal CSF composition (**Figure 6B**).

**Figure 6.**
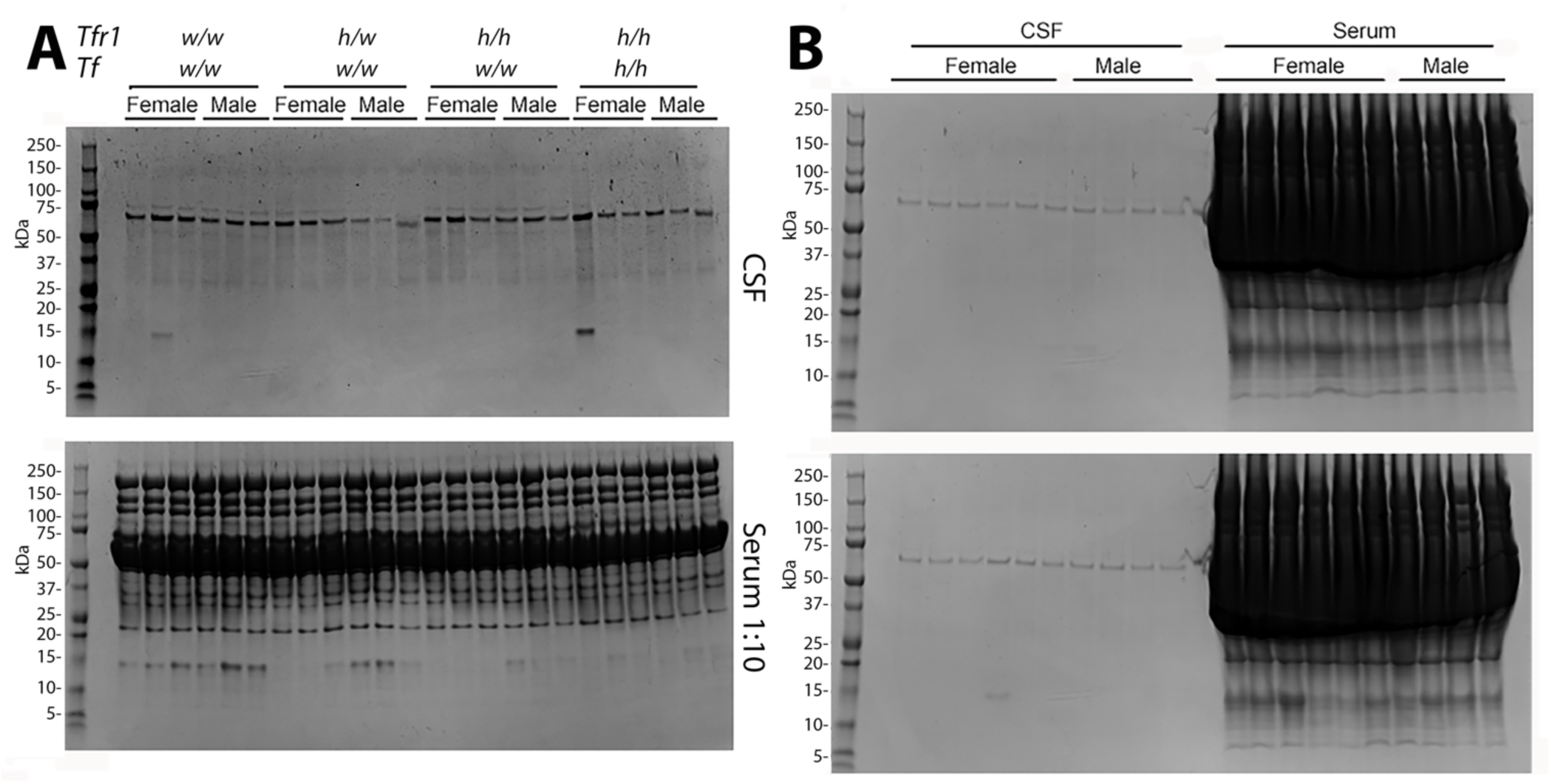
The BBB remains intact in humanized TfR1 rats. **A)** SDS-PAGE followed by Coomassie Blue staining of equal volumes of serum and CSF) from rats with humanized alleles at the *Tfr1* and *Tf* gene loci. The protein content and pattern in CSF are similar between control rats (*Tf^w/w^:Tfr1^w/w^*) and those carrying heterozygous or homozygous humanized *TfR1* alleles (*Tf^w/w^:Tfr1^h/w^* and *Tf^w/w^:Tfr1^h/h^*), as well as rats homozygous for both human *Tf* and *Tfr1* (*Tf^h/h^:Tfr1^h/h^*), indicating that BBB integrity is preserved following humanization. A ∼15 kDa band of unknown identity is visible in the CSF of one control (*Tf^w/w^:Tfr1^w/w^*) and one double-humanized rat(*Tf^h/h^:Tfr1^h/h^*). Serum samples (diluted 1:10 in PBS prior to loading) show comparable protein patterns across genotypes. **B)** Additional *Tfr1^h/h^* rat samples confirm the findings. While serum lanes display abundant protein bands, only a faint ∼70 kDa band is detected in CSF, consistent with the low protein content of normal CSF and exclusion of serum proteins by an intact BBB.

Thus, we used *Tfr1^h^* rats to assess whether TfR1b-Nbs from Families A, B, D, and G can cross the BBB. **Figure 7** summarizes the results of the in vivo brain permeability experiment, with each dot representing an individual animal. The data shown in **Figure 7A** are compiled from all animals included in the experiment detailed in **Table 2**. As shown, all tested TfR1b-Nbs were detected in brain homogenates of *Tfr1^h/h^* and *Tfr1^h/w^* rats, whereas no signal was observed in animals expressing only the endogenous rat *Tfr1* (*Tfr1^w/w^*). These results support the conclusion that TfR1b-Nbs can mediate transcytosis across the BBB via human TfR1, and that the presence of even a single human *TfR1* allele is sufficient to enable BBB crossing. Notably, serum levels of the nanobodies were also markedly higher in *Tfr1^h/h^* and *Tfr1^h/w^* rats, likely reflecting increased stability due to target engagement. The data were pooled because the number of animals injected with each individual nanobody was too small to allow for meaningful analysis or visualization on a per-nanobody basis. Additionally, heterogeneity in sex, age, and genotype across experimental groups further complicated presentation of the results for each nanobody individually. Nevertheless, the aggregated dataset in **Figure 7** clearly demonstrates that targeting human TfR1 with nanobodies enables TfR1-dependent transcytosis into the brain. It should be noted, however, that this pooled representation does not permit a comparative evaluation of the relative efficiency of each nanobody.

**Figure 7.**
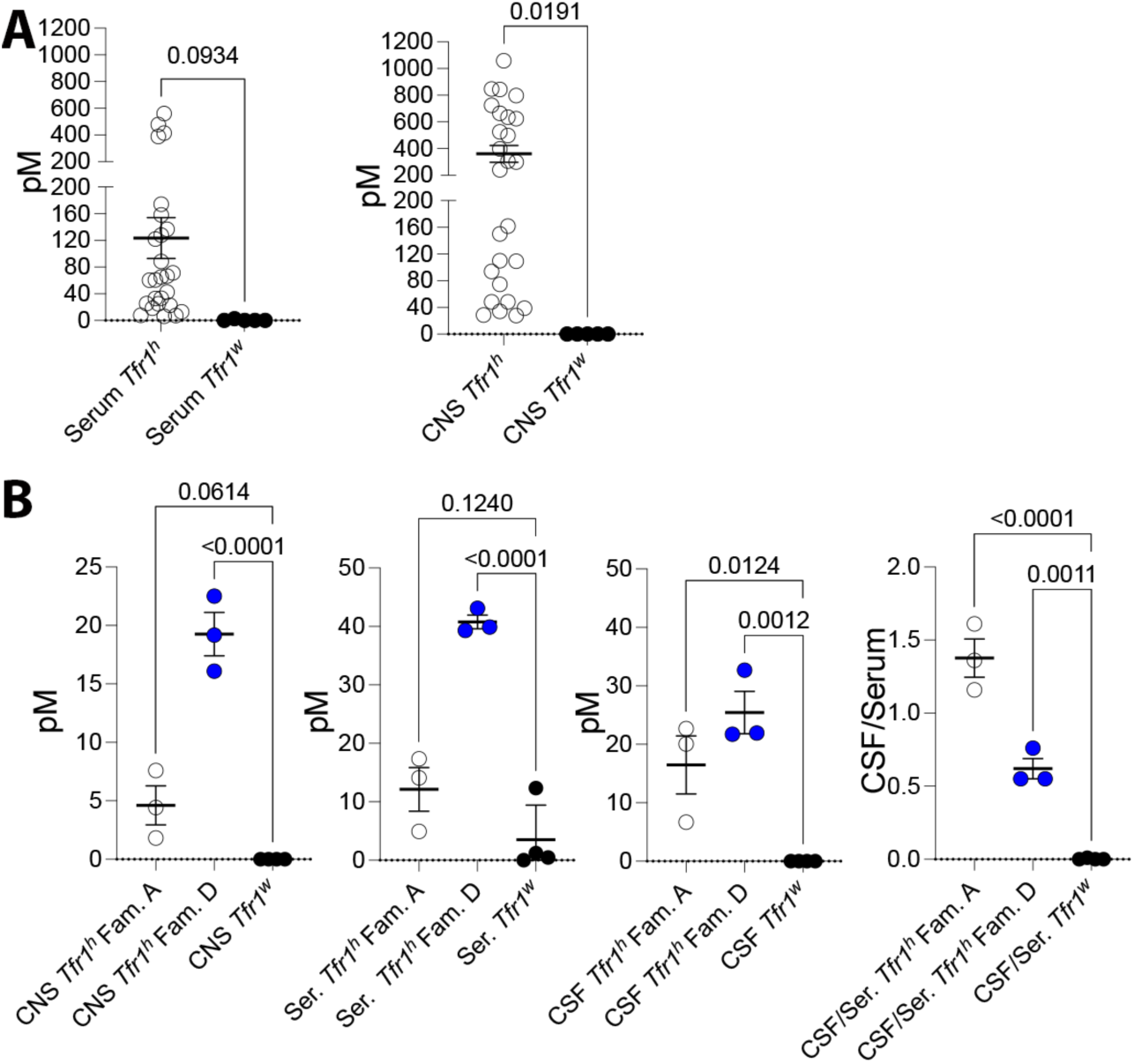
In vivo BBB permeability of TfR1b-Nbs in humanized Tfr1 rats. **A)** ELISA detection of TfR1b-Nbs in brain homogenates of rats expressing human TfR1 (*Tfr1^h/w^* or *Tfr1^h/h^*), but not in wild-type rats (*Tfr1^w/w^*), 16-18 hrs. following IV injection. **B)** Detection of TfR1b-Nbs 04B05R3 (Family A) and 05D01R3 (Family D) in serum, CSF, and brain homogenates of *Tfr1^h/w^* and *Tfr1^w/w^* rats 44–48 hours post-injection. Both nanobodies showed high CSF/serum ratios in *Tfr1^h/w^* rats, indicating efficient BBB penetration and sustained stability.

**Table 2.** Detection of TfR1b-Nbs in brain samples of rats expressing human TfR1, compared with rat Tfr1-expressing animals. Columns 1–6 list the nanobody name, family, *Tfr1* genotype, sex, age, and time of tissue collection post-injection. Columns 7 and 8 report the concentrations of TfR1b-Nbs in serum and brain homogenates, respectively. Data for this experiment are shown in Figure 6A.

| Nanobody | Family | *Tfr1* | sex | Age in days | Time (hrs) | Conc. in Serum [pM] | Conc. in Brain homog. [pM] |
| --- | --- | --- | --- | --- | --- | --- | --- |
| 02B02R3 | A | *h/h* | f | 70 | 16-18 | 33.2 | 28.6 |
| 05F02R3 | A | *h/h* | f | 70 | 16-18 | 22.7 | 28.0 |
| 04F01R3 | B | *h/h* | m | 39 | 16-18 | 128.1 | 241.2 |
| 04G05R3 | D | *h/h* | m | 39 | 16-18 | 414.8 | 624.4 |
| 03E01R3 | G | *h/h* | m | 31 | 16-18 | 559.7 | 843.8 |
| 04E01R3 | G | *h/h* | f | 34 | 16-18 | 388.2 | 796.9 |
| 01B01R3 | I | *h/h* | m | 122 | 16-18 | 60.6 | 109.9 |
| 01B01R3 | I | *h/h* | f | 34 | 16-18 | 60.4 | 299.0 |
| 03G10R3 | I | *h/h* | m | 121 | 16-18 | 13.0 | 48.4 |
| 02B02R3 | A | *h/w* | m | 114 | 16-18 | 42.4 | 94.0 |
| 02F06R3 | A | *h/w* | m | 61 | 16-18 | 71.2 | 161.8 |
| 04B05R3 | A | *h/w* | f | 107 | 12 | 478.5 | 1058.6 |
| 04B05R3 | A | *h/w* | f | 237 | 16-18 | 174.1 | 525.2 |
| 05D05R3 | A | *h/w* | m | 237 | 16-18 | 25.0 | 39.0 |
| 05F02R3 | A | *h/w* | f | 147 | 16-18 | 33.4 | 109.3 |
| 04D08R3 | B | *h/w* | m | 147 | 16-18 | 66.7 | 498.6 |
| 04D08R3 | B | *h/w* | m | 34 | 16-18 | 25.4 | 48.2 |
| 04F01R3 | B | *h/w* | m | 114 | 16-18 | 88.5 | 635.1 |
| 04G05R3 | D | *h/w* | m | 147 | 16-18 | 121.9 | 847.2 |
| 03E01R3 | G | *h/w* | f | 64 | 16-18 | 158.3 | 663.6 |
| 04E01R3 | G | *h/w* | f | 212 | 16-18 | 136.6 | 724.3 |
| 05D01R3 | G | *h/w* | f | 94 | 12 | 65.3 | 398.1 |
| 05D01R3 | G | *h/w* | m | 61 | 16-18 | 18.6 | 306.8 |
| 01B01R3 | I | *h/w* | m | 149 | 16-18 | 7.6 | 74.5 |
| 01B06R3 | I | *h/w* | m | 65 | 16-18 | 6.3 | 34.1 |
| 03G10R3 | I | *h/w* | m | 51 | 16-18 | 7.8 | 150.1 |
| 02F06R3 | A | *w/w* | m | 49 | 16-18 | 0.0 | 0.0 |
| 04B05R3 | A | *w/w* | f | 217 | 16-18 | 0.0 | 0.0 |
| 05D05R3 | A | *w/w* | m | 217 | 16-18 | 2.8 | 0.0 |
| 05D01R3 | G | *w/w* | m | 51 | 16-18 | 0.0 | 0.0 |
| 04G05R3 | D | *w/w* | f | 139 | 12 | 0.0 | 0.0 |

In a follow-up experiment, nanobodies 04B05R3 (Family A) and 05D01R3 (Family G) were tested again in young *Tfr1^h/w^* or control *Tfr1^w/w^* rats. **Table 3** shows detailed information about age, sex, genotype of the rats utilized and nanobodies injected. Samples were collected 44–48 hours post-IV injection, including CSF. As before, serum levels were higher in *Tfr1^h/w^* rats, and only these rats showed TfR1b-Nbs in both the brain and CSF (**Figure 7B**). Both nanobodies exhibited remarkably high CSF/serum ratios, with 04B05R3 reaching ∼1.5, suggesting preferential brain accumulation. These nanobodies remained detectable 2 days post-injection, reflecting prolonged stability—likely enhanced by target engagement—an uncommon feature for nanobodies, which typically have short half-lives.

**Table 3.** Detection of TfR1b-Nbs in brain samples of rats expressing human TfR1, compared with rat Tfr1-expressing animals. A human TfR1-specific ELISA detected TfR1b-Nbs in brain tissues and CSF of *Tfr1^h/w^*rats, but not in *Tfr1^w/w^* controls. Columns 1–6 list the nanobody name, family, *Tfr1* genotype, sex, age, and time of tissue collection post-injection. Columns 7–10 report the concentrations of TfR1b-Nbs in serum, CSF, the CSF/serum ratio, and brain homogenates, respectively. Data for this experiment are shown in Figure 6B.

| Nanobody | Family | *Tfr1* | sex | Age in days | Time (hrs) | Serum [pM] | CSF[pM] | CSF/Serum | CNS [pM] |
| --- | --- | --- | --- | --- | --- | --- | --- | --- | --- |
| 04B05R3 | A | *h/w* | m | 43 | 44-48 | 14.101 | 22.706 | 1.61 | 4.431 |
| 04B05R3 | A | *h/w* | m | 43 | 44-48 | 17.371 | 20.066 | 1.16 | 7.593 |
| 04B05R3 | A | *h/w* | f | 42 | 44-48 | 4.927 | 6.6769 | 1.36 | 1.83 |
| 05D01R3 | G | *h/w* | m | 44 | 44-48 | 43.093 | 32.691 | 0.76 | 22.517 |
| 05D01R3 | G | *h/w* | m | 44 | 44-48 | 39.896 | 21.941 | 0.55 | 19.183 |
| 05D01R3 | G | *h/w* | f | 44 | 44-48 | 39.326 | 21.714 | 0.55 | 16.104 |
| 04B05R3 | A | *w/w* | m | 42 | 44-48 | 1.226 | 0 | 0 | 0 |
| 04B05R3 | A | *w/w* | m | 44 | 44-48 | 0.498 | 0 | 0 | 0 |
| 05D01R3 | G | *w/w* | m | 42 | 44-48 | 0 | 0 | 0 | 0 |
| 05D01R3 | G | *w/w* | m | 43 | 44-48 | 12.371 | 0 | 0 | 0 |

### Assessing CNS delivery of BBB-impermeable biologics via TfR1b-Nb-directed transcytosis

We have also produced nanobodies that inhibit TNFα activity, referred to as TNFI-Nbs. **Figure 8A** shows that TNFI-Nb was detected in the serum of both control (*Tfr1^w/w^*) and humanized (*Tfr1^h/w^*) rats following intravenous injection (see **Table 4** for experimental details). However, it was undetectable in brain homogenates of either genotype, indicating that TNFI-Nb does not cross the BBB. This absence confirms that the humanized BBB remains functionally restrictive to non-transported proteins.

**Figure 8.**
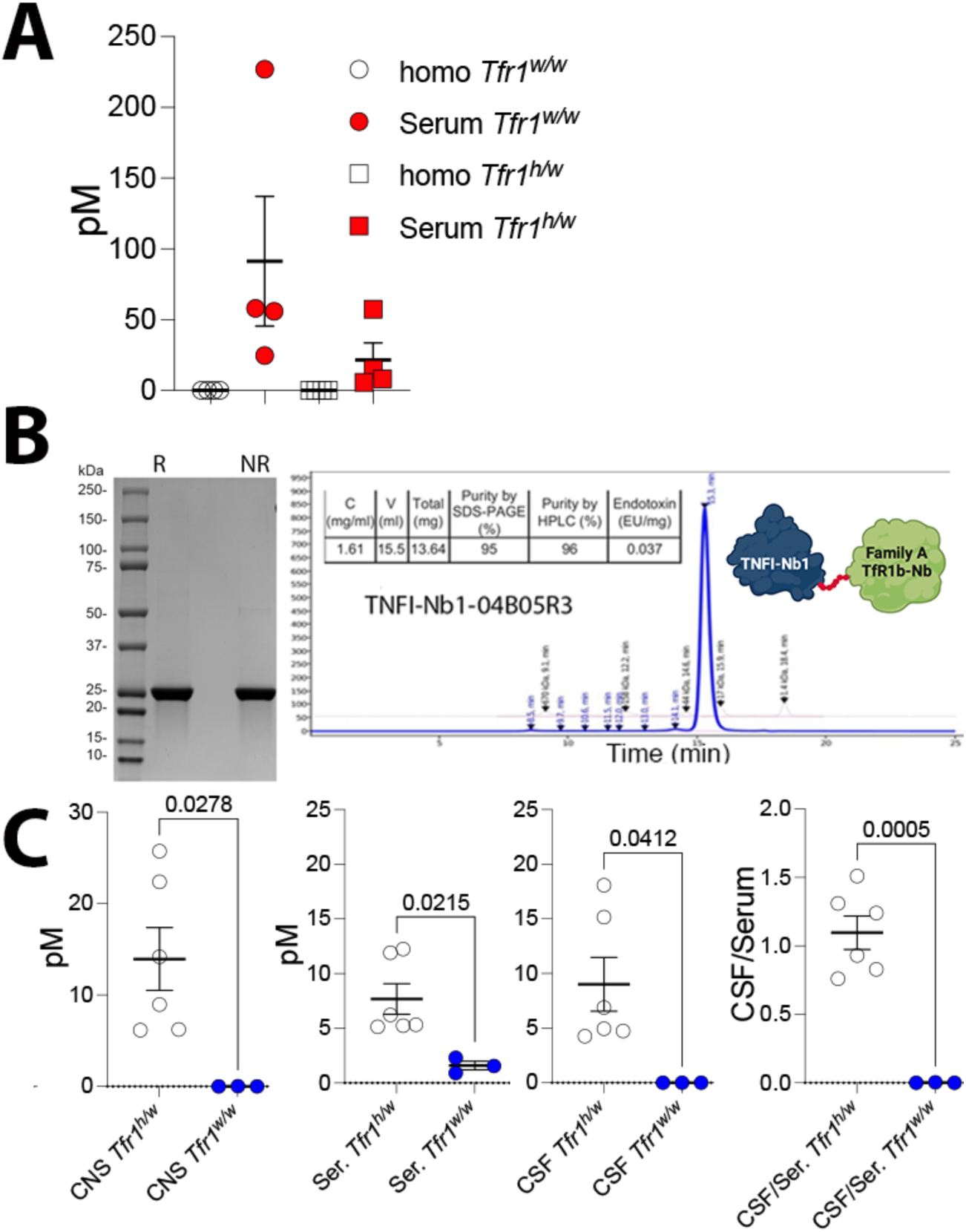
Fusion to TfR1b-Nb enables CNS delivery of TNFI-Nb1 via human TfR1-mediated transcytosis. **A)** TNFα-based ELISA quantification of TNFI-Nb1 in serum and brain homogenates from *Tfr1^h/w^* and *Tfr1^w/w^* rats, 6 hours after intravenous injection. TNFI-Nb1 was detectable in serum but absent in brain tissue, indicating lack of BBB permeability. **B)** SDS-PAGE and SEC-HPLC analysis of the heterodimer composed of TNFI-Nb1 fused to TfR1b-Nb 04B05R3 (Family A), produced in CHO-S cells. **C**) TNFα-based ELISA quantification of TNFI-Nb1∼04B05R3 in serum, CSF, and brain homogenates of *Tfr1^h/w^*and *Tfr1^w/w^* rats, 48 hours post-injection. The heterodimer was present in serum of both genotypes but detected in CSF and brain only in *Tfr1^h/w^* rats, demonstrating human TfR1-dependent BBB transcytosis. The observed CSF/serum ratio of ∼1 indicates efficient CNS delivery.

**Table 4.** Details of the animals used in the experiment presented in Figure 7, including genotype, sex, age, and nanobody administered.

| Nanobody | *Tfr1* | sex | Age in days | Time (hrs) | Serum [pM] | CNS [pM] |
| --- | --- | --- | --- | --- | --- | --- |
| TNFI-Nb1 | *h/w* | f | 48 | 6 | 14.81634 | 0 |
| TNFI-Nb1 | *h/w* | f | 48 | 6 | 5.659766 | 0 |
| TNFI-Nb1 | *h/w* | m | 40 | 6 | 8.185005 | 0 |
| TNFI-Nb1 | *h/w* | f | 108 | 6 | 57.38742 | 0 |
| TNFI-Nb1 | *w/w* | f | 139 | 6 | 226.9179 | 0 |
| TNFI-Nb1 | *w/w* | f | 39 | 6 | 55.92783 | 0 |
| TNFI-Nb1 | *w/w* | m | 39 | 6 | 57.97242 | 0 |
| TNFI-Nb1 | *w/w* | m | 39 | 6 | 24.78752 | 0 |

Next, we evaluated whether fusing TNFI-Nb1 to the TfR1b-Nb 04B05R3 (Family A) could enhance brain delivery of TNFI-Nb1. The heterodimer was produced in CHO-S cells by GenScript (**Figure 8B**). The TNFI-Nb1 and TfR1b-Nb 04B05R3 nanobodies were genetically fused using a flexible glycine-serine–rich linker with the sequence GGGSGGGSGGGSGGG. This linker was selected based on two key criteria: 1) It provides structural flexibility while minimizing the risk of steric interference between the two domains; 2) It has low predicted immunogenicity, as it is not expected to generate peptides capable of stimulating T cell responses, thereby reducing the likelihood of anti-drug immune reactions.

To assess CNS transcytosis, we employed a TNFα-specific ELISA. The same protocol used for monomeric TfR1b-Nbs was applied, including IV injection, perfusion, and sample collection. **Table 5** summarizes the age, sex, genotype of the rats used, and the nanobodies injected. The heterodimer was detected in both the brain and CSF of *Tfr1^h/w^* rats, but not in or control *Tfr1^w/w^* controls, confirming human TfR1-dependent BBB transport (**Figure 8C**). Like the monomeric 04B05R3, the heterodimer exhibited strong CNS localization, with a CSF/serum ratio of ∼1—slightly lower than the monomer (∼1.5) yet still indicating efficient brain accumulation.

**Table 5.** Shuttle Activity of TfR1b-Nbs for TNFI-Nb1 in brain samples of rats expressing human TfR1. A human TNFα-specific ELISA detected TNFI-Nb1-04B05R3 in brain tissues and CSF of *Tfr1^h/w^*rats, but not in *Tfr1^w/w^* controls. The linker used to link these two nanobodies was: GGGGSGGG. Columns 1–6 list the nanobody name, family, *Tfr1* genotype, sex, age, and time of tissue collection post-injection. Columns 7–10 report the concentrations of TfR1b-Nbs in serum, CSF, the CSF/serum ratio, and brain homogenates, respectively. Data for this experiment are shown in Figure 8.

| Nanobody | Family | *Tfr1* | sex | Age in days | Time (hrs) | Serum [pM] | CSF[pM] | CSF/Serum | CNS [pM] |
| --- | --- | --- | --- | --- | --- | --- | --- | --- | --- |
| TNFI-Nb1-04B05R3 | A | *h/w* | m | 44 | 44-48 | 5.14 | 4.25 | 0.83 | 6.23 |
| TNFI-Nb1-04B05R3 | A | *h/w* | f | 44 | 44-48 | 5.32 | 4.95 | 0.93 | 6.17 |
| TNFI-Nb1-04B05R3 | A | *h/w* | m | 44 | 44-48 | 6.21 | 4.73 | 0.76 | 8.96 |
| TNFI-Nb1-04B05R3 | A | *h/w* | f | 51 | 44-48 | 5.24 | 6.88 | 1.31 | 14.18 |
| TNFI-Nb1-04B05R3 | A | *h/w* | m | 50 | 44-48 | 12.23 | 15.16 | 1.24 | 25.73 |
| TNFI-Nb1-04B05R3 | A | *h/w* | m | 51 | 44-48 | 11.91 | 18.08 | 1.51 | 22.40 |
| TNFI-Nb1-04B05R3 | A | *w/w* | m | 51 | 44-48 | 0.91 | 0.004 | 0.004 | 0.00 |
| TNFI-Nb1-04B05R3 | A | *w/w* | f | 44 | 44-48 | 2.30 | 0.004 | 0.001 | 0.03 |
| TNFI-Nb1-04B05R3 | A | *w/w* | f | 44 | 44-48 | 1.56 | 0.002 | 0.001 | 0.00 |

To further evaluate BBB transcytosis of the heterodimer, we applied a fractionation protocol (Kariolis et al., 2020) used by Denali to assess CNS penetration of AVI-based therapeutics. One hemibrain from each of three *Tfr1^h/w^* rats was separated into vasculature and vascular-depleted parenchymal fractions. In this experiment, we tested the TNFI-Nb1– linker–04B05R3 heterodimer.

Western blot analysis confirmed effective fractionation: the endothelial marker Glut1 was enriched in the vasculature fraction and depleted in the parenchyma, while Gapdh—a general cellular marker—was enriched in the parenchyma. Cell-type–specific markers in the parenchymal fraction included: Vamp2 (neurons), NmdaR2b (neurons), Iba1(microglia), Eaat2 (astrocytes), and Mbp (oligodendrocytes) (**Figure 9A**). These results indicate successful enrichment of parenchymal CNS cell types, with particularly strong signal for microglial marker. Human TfR1 was detected in all brain fractions of *Tfr1^h/w^* rats, with highest expression in the vasculature, consistent with its known highest endothelial localization.

**Figure 9.**
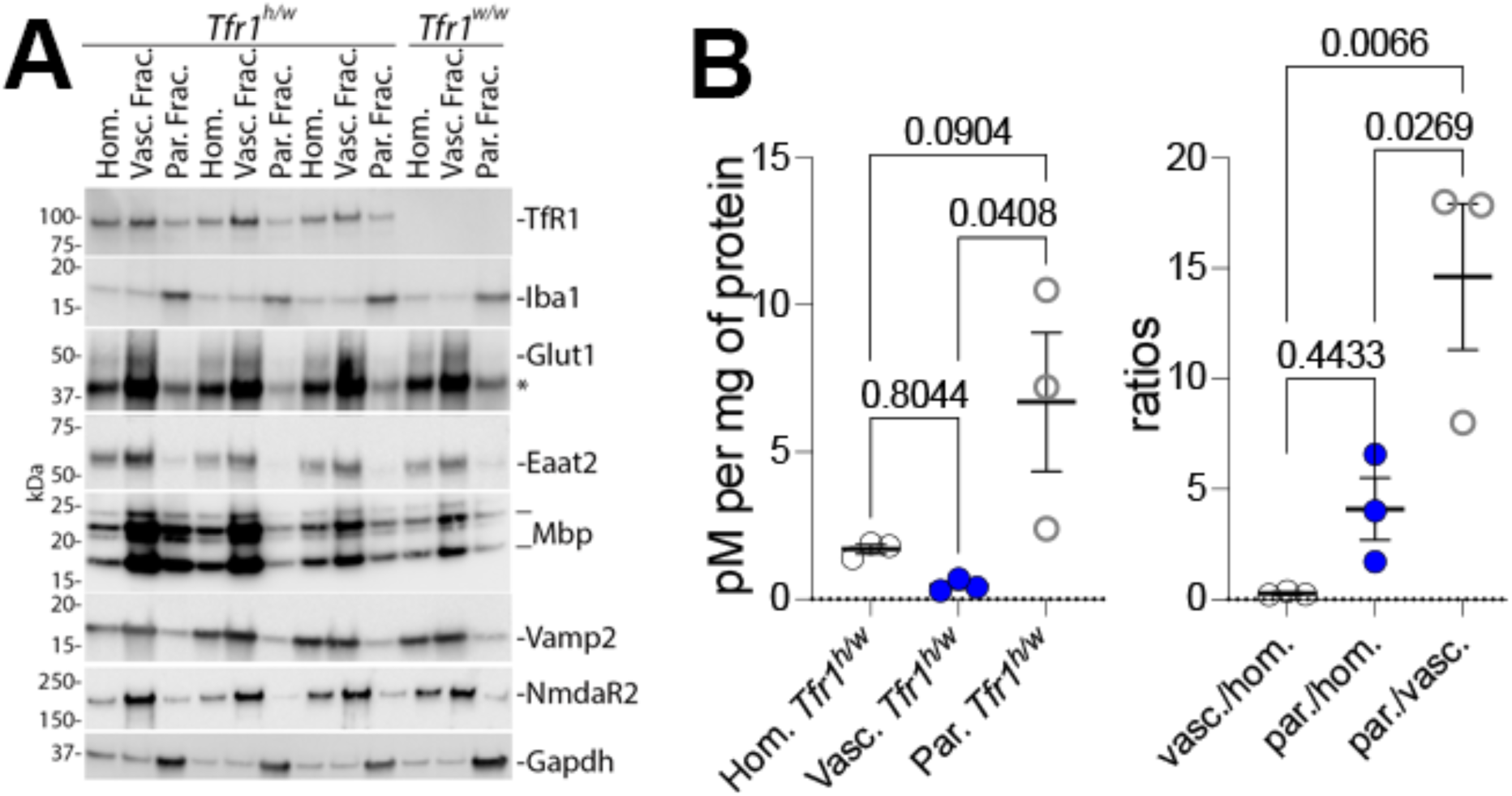
Brain fractionation analysis of TNFI-Nb1–linker–04B05R3 distribution in Tfr1^h/w^ rats. **A)** Western blot validation of brain fractionation into vasculature and parenchyma. Glut1 (endothelial marker) was enriched in the vasculature fraction, while Gapdh and cell-type–specific markers—Vamp2 and NmdaR2b (neurons), Iba1 (microglia), Eaat2 (astrocytes), and Mbp (oligodendrocytes)—were enriched in the parenchymal fraction. Human TfR1 was detected in all fractions, with strongest signal in the vasculature. **B)** ELISA quantification of TNFI-Nb1–04B05R3 in homogenate, vasculature, and parenchymal fractions. After normalization to protein content, the parenchymal fraction showed the highest nanobody levels, supporting uptake by CNS cells via human TfR1-mediated transcytosis.

ELISA analysis (**Figure 9B**) confirmed the presence of TNFI-Nb1–04B05R3 in brain homogenates, vasculature, and parenchymal fractions of *Tfr1^h/w^* rats. After normalization to protein content, the parenchymal fraction showed the highest levels, supporting uptake of the heterodimer by CNS cells, likely via human TfR1.

These results support the concept that TfR1b-Nbs can serve as effective “NewroBuses,” delivering otherwise BBB-impermeable biologics into the brain.

### Assessment of NewroBus-directed brain uptake of a non–BBB-penetrant biologic

To confirm human TfR1-dependent BBB penetration of TNFI-Nb1-04B05R3, we performed immunohistochemistry (IHC). One hemibrain from each rat used in the experiment shown in **Figure 9** was fixed in 4% paraformaldehyde following perfusion. Brain sections from *Tfr1^h/w^* and *Tfr1^w/w^* rats were stained using anti-His tag (for the nanobody) and anti-human TfR1 antibodies to examine nanobody localization and target engagement. TNFI-Nb1–04B05R3 (green) was detected in *Tfr1^h/w^*but not *Tfr1^w/w^* brains (**Figure 10A**). **Figure 10B** confirms human TfR1 expression (red) exclusively in *Tfr1^h/w^* brains. Colocalization of TNFI-Nb1–04B05R3 (green), human TfR1 (red), and the endothelial marker CD31 (purple) was observed in *Tfr1^h/w^* rats (**Figure 10C**). Merged and zoomed-in images show clear colocalization (white), confirming binding to the brain endothelium in a TfR1-dependent manner.

**Figure 10.**
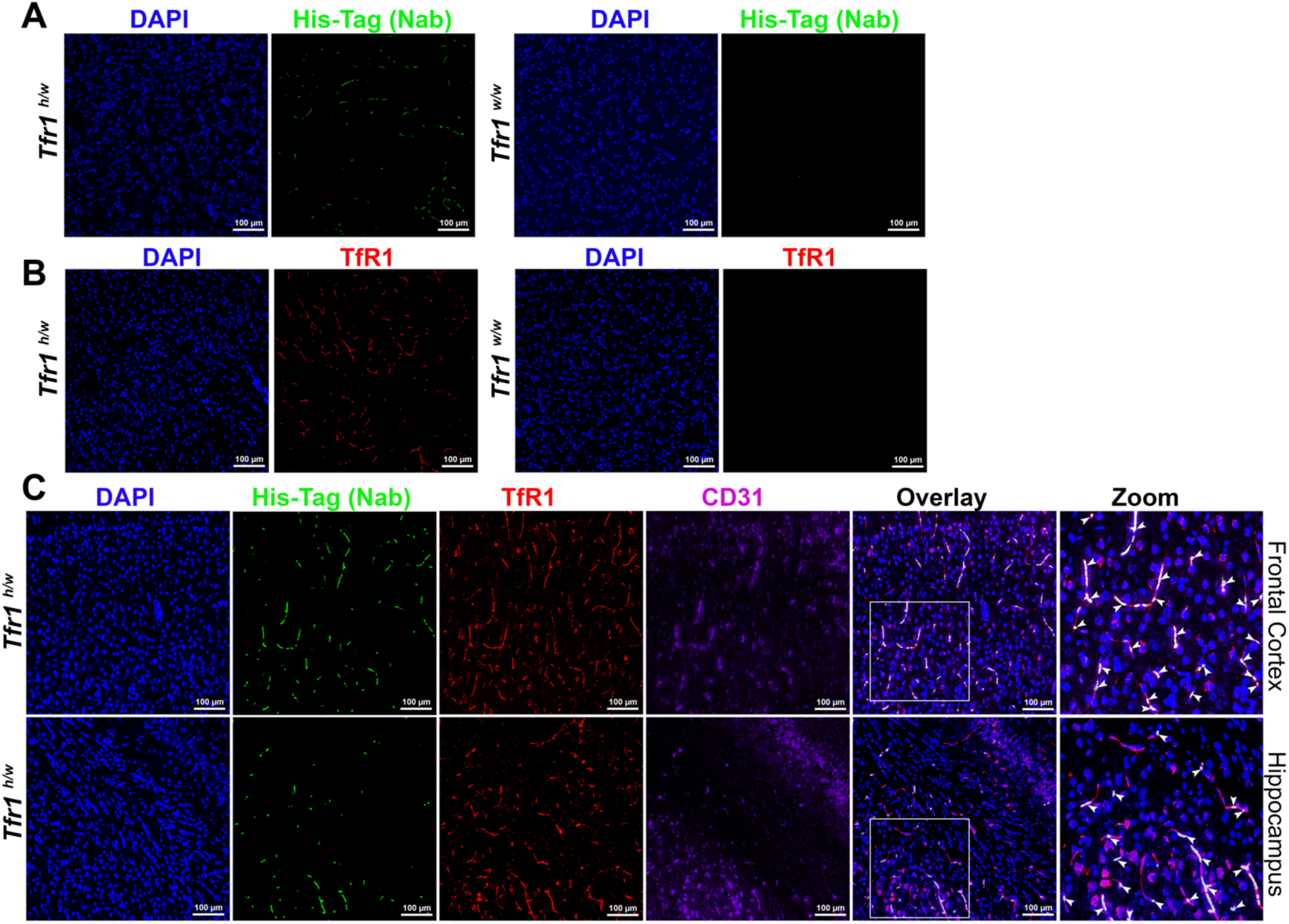
TNFI-Nb1-linker-04B05R3 colocalize with human TfR1in brain vessels of *Tfr1^h/w^* rats. **A**) Anti-His tag antibody (green) and (**B**) anti-human TfR1 antibody (red) stain TNFI-Nb1-linker-04B05R3 and human TfR1, respectively, in the brains of *Tfr1^h/w^* but not *Tfr1^w/w^* rats. **C**) Human TfR1 (red), CD31 (purple)—an endothelial cell marker—and TNFI-Nb1-linker-04B05R3 (green) colocalize in vessels in the brains of *Tfr1^h/w^* rats. White arrows in the zoomed-in panel indicate the colocalization spots within the region highlighted by the white square in the overlay panel.

The presence of the heterodimer in the parenchymal fraction suggests cellular uptake within the CNS. To investigate whether TNFI-Nb1–04B05R3 localizes to specific brain cell types, we performed confocal imaging of cortex and hippocampus sections stained with DAPI (nuclei), GFAP (astrocytes), IBA1 (microglia), and an anti-His tag antibody to detect TNFI-Nb1–04B05R3. TNFI-Nb1–04B05R3 colocalizes with GFAP-positive astrocytes (yellow signal) in both cortex and hippocampus of *Tfr1^h/w^* rats, but not in *Tfr1^w/w^*controls (**Figure 11**). **Figure 12** shows similar colocalization with IBA1-positive microglia, again only in *Tfr1^h/w^*rats.

**Figure 11.**
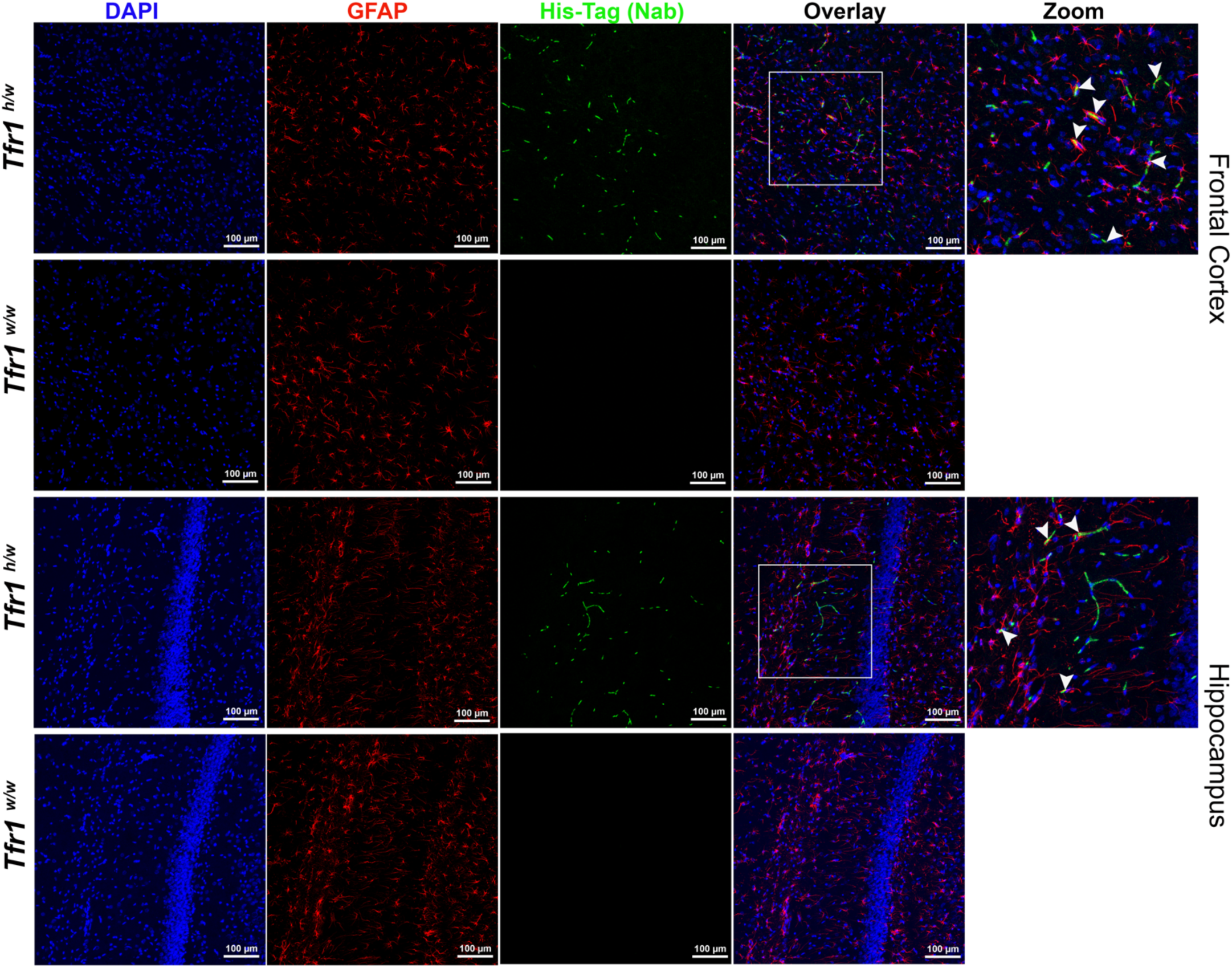
Confocal images showing the localization of TNFI-Nb1-linker-04B05R3 in astrocytes of the cortex and hippocampus in *Tfr1^h/w^* and *Tfr1^w/w^* rats. Sections were stained with an anti-His tag antibody (green) to detect TNFI-Nb1-linker-04B05R3 and an anti-GFAP antibody (red) to label astrocytes. TNFI-Nb1-linker-04B05R3 is present in the brains of *Tfr1^h/w^* rats but not in *Tfr1^w/w^* rats. In *Tfr1^h/w^* rats, TNFI-Nb1-linker-04B05R3 colocalizes with some astrocytes, indicated by yellow overlays in the merged images. White arrows in the zoomed-in panel point to colocalization spots within the region highlighted by the white square in the overlay panel.

**Figure 12.**
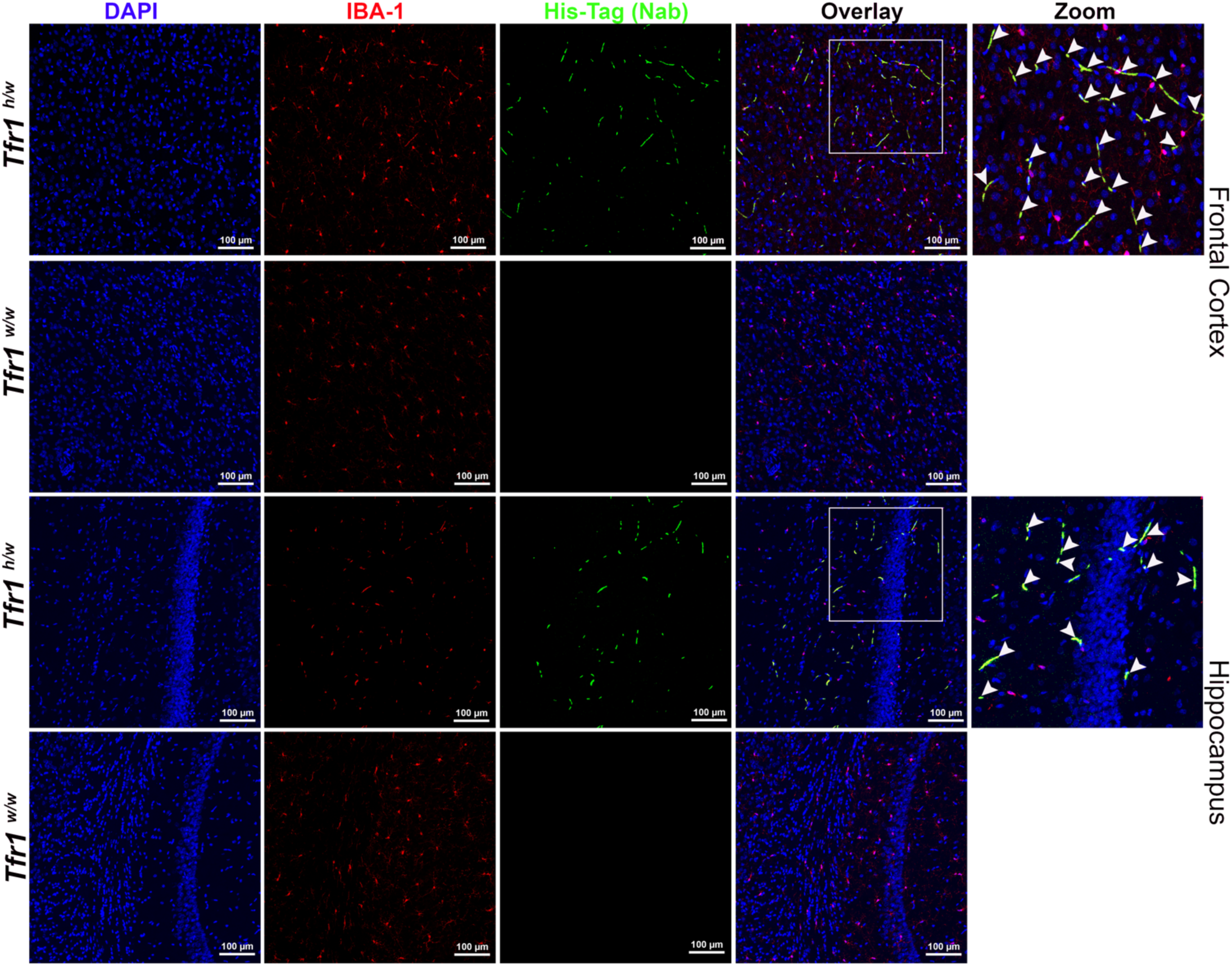
Confocal images showing the localization of TNFI-Nb1-linker-04B05R3 in microglia of the cortex and hippocampus in *Tfr1^h/w^* and *Tfr1^w/w^* rats. Sections were stained with an anti-His tag antibody (green) to detect TNFI-Nb1-linker-04B05R3 and an anti-IBA1 antibody (red) to label microglia. TNFI-Nb1-linker-04B05R3 is present in the brains of *Tfr1^h/w^* rats but not in *Tfr1^w/w^* rats. In *Tfr1^h/w^* rats, TNFI-Nb1-linker-04B05R3 colocalizes with most microglia, as indicated by yellow overlays in the merged images. White arrows in the zoomed-in panel point to colocalization spots within the region highlighted by the white square in the overlay panel.

Together, these data demonstrate that TNFI-Nb1–04B05R3 crosses the BBB in a human TfR1-dependent manner, localizing to the CSF—an accessible surrogate for interstitial fluid within the brain parenchyma—with a portion taken up by CNS cells, predominantly astrocytes and microglia. The absence of signal in *Tfr1^w/w^* rats underscores the requirement for human TfR1. These findings support the potential of TfR1b-Nbs as targeted delivery vehicles for otherwise BBB-impermeable therapeutics.

### In silico humanization and developability optimization of lead TfR1b nanobodies

Humanization and optimization are critical steps to enhance nanobody stability, solubility, and expression, while minimizing immunogenicity—key for safe and effective therapeutic use in humans. Accordingly, TfR1b-Nbs from Families A, B, D, and G underwent in silico humanization and developability optimization using a strategy similar to that applied to TNFI-Nbs ^11^. We employed AbNatiV, a machine learning algorithm that predicts both humanness and VHH-nativeness directly from sequence ^12^. Humanness scores above 0.8 indicate high similarity to human variable domains and correlate with lower immunogenicity risk, while scores below 0.8 suggest greater likelihood of immune recognition due to non-human sequence features.

Based on the most favorable starting points for humanness, VHH-nativeness, and CamSol solubility ^13^, TFRI1b-Nbs 05F02, 04F01, and 04G05 were selected for TNF1b-Nbs Families A, B, and D, respectively. For Family G, 03E01 and 05D01 were chosen as representatives of the two subfamilies present in this group (**Table 6**).Two complementary mutational sampling strategies were applied: enhanced sampling, which rapidly converges on an optimized humanized sequence, and exhaustive sampling, which systematically evaluates all permissible residue substitutions based on position-specific scoring matrices (PSSMs) from human VH and VHH databases. Both methods identify designs on the Pareto front, balancing increased humanness with preserved VHH-nativeness—critical for maintaining nanobody stability and folding in the absence of a VL domain.

**Table 6.** AbNatiV Assessment of TfR1b-Nbs. Key metrics include Humaness (H), VHH-ness (VH), and CamSol Intrinsic (S). The Nbs underlined, in bold and italic have been selected for humanization.

| ID | F | H | V | S |
| --- | --- | --- | --- | --- |
| 02F06 | **A** | 0.721 | 0.744 | 0.382 |
| 05D05 |  | 0.697 | 0.783 | 0.405 |
| ***<u>05F02</u>*** |  | 0.754 | 0.787 | 0.357 |
| 02B02 |  | 0.745 | 0.785 | 0.291 |
| 04B05 |  | 0.734 | 0.772 | 0.304 |
| 04D08 | **B** | 0.774 | 0.831 | 0.620 |
| ***<u>04F01</u>*** |  | 0.763 | 0.847 | 0.628 |
| ***<u>04G05</u>*** | **D** | 0.763 | 0.850 | 0.438 |
| ***<u>05D01</u>*** | **G** | 0.634 | 0.732 | 0.505 |
| ***<u>03E01</u>*** |  | 0.619 | 0.719 | 0.042 |
| 04E01 |  | 0.601 | 0.735 | 0.026 |

Humanization was restricted to solvent-exposed framework residues, while CDRs were preserved to avoid compromising antigen binding. As a further control for structural integrity post-humanization, the structures of wild-type (WT) and all humanized sequences were modelled with NanobodyBuilder 2 ^14^. The modelled structures were superimposed based on their framework regions, and the root-mean-square deviation (RMSD) calculated for the CDR regions was approximately 1 Å. This value is significantly smaller than the expected modelling accuracy for these regions suggesting minimal or no displacement of the CDR loops resulting from the humanizing framework mutations. Following in silico humanization, we used the structural models of both Enhanced and Exhaustive humanized variants as inputs for the CamSol Combination pipeline ^15^, by excluding all CDR regions from the design and by using an alignment of human VH sequences as input, rather than the default VHH sequences. CamSol Combination automatically identifies combinations of mutations predicted to improve solubility and conformational stability, or one of these properties without affecting the other. As a further computational filter, the apparent melting temperature of in silico mutants was predicted with NanoMelt ^16^. Of the mutations suggested by CamSol Combination, we retained only those that didn’t reduce humanness according to AbNatiV scoring (**Table 7**). These variants were produced in CHO-S cells with production performed at GenScript.

**Table 7.** TfR1b-Nb mutants with enhanced therapeutic potential determined by AbNatiV analysis. Changes in Humanness (H), VHH-ness (VH), and CamSol Intrinsic (S) are presented in the last three columns.

| ID | H | V | S | Code | Fam. |
| --- | --- | --- | --- | --- | --- |
| **05F02** | 0.754 | 0.787 | 0.357 |  | **A** |
| Enhanced WT-CDR3STEMS | 0.790 | 0.751 | 0.405 |  |  |
| enhancedWT-CDR3STEM-W | 0.828 | 0.768 | 0.299 | **A2** |  |
| enhanced | 0.841 | 0.782 | 0.504 | **A3** |  |
| Enhanced WT-CDR3STEM-W v1 | 0.821 | 0.769 | 0.598 |  |  |
| enhanced v3 | 0.840 | 0.753 | 0.602 |  |  |
| exhaustive 7 | 0.868 | 0.812 | 0.441 | **A4** |  |
| enhanced v1 | 0.812 | 0.749 | 0.800 |  |  |
| enhanced v2 | 0.827 | 0.769 | 0.802 |  |  |
| **04F01** | 0.763 | 0.847 | 0.628 |  | **B** |
| Enhanced WT-CDR3STEM | 0.870 | 0.869 | 0.828 | **B1** |  |
| enhanced | 0.882 | 0.866 | 0.960 |  |  |
| Enhanced WT-CDR3STEM v1 | 0.867 | 0.878 | 0.823 | **B2** |  |
| enhanced v1 | 0.884 | 0.850 | 1.029 |  |  |
| exhaustive 12 | 0.899 | 0.876 | 0.978 |  |  |
| enhanced v2 | 0.882 | 0.869 | 0.991 |  |  |
| enhanced v3 | 0.884 | 0.858 | 1.023 |  |  |
| **04G05** | 0.763 | 0.850 | 0.438 |  | **D** |
| Enhanced WT-CDR3Stem | 0.846 | 0.860 | 0.632 | **D1** |  |
| enhanced | 0.854 | 0.863 | 0.847 |  |  |
| Enhanced WT-CDR3Stem v1 | 0.834 | 0.854 | 0.667 | **D2** |  |
| exhaustive | 0.843 | 0.841 | 0.572 | **D3** |  |
| enhanced v1 | 0.814 | 0.838 | 0.939 |  |  |
| enhanced v2 | 0.831 | 0.854 | 0.888 |  |  |
| **05D01** | 0.763 | 0.850 | 0.438 |  | **G** |
| enhanced QAP insertion | 0.802 | 0.841 | 0.641 |  |  |
| enhanced QAP insertion v1 | 0.796 | 0.833 | 0.711 |  |  |
| **03E01** | 0.619 | 0.719 | 0.042 |  |  |
| exhaustive QAP insertion | 0.793 | 0.817 | 0.584 |  |  |
| enhanced | 0.784 | 0.820 | 0.500 |  |  |

CamSol Combination automatically identifies combinations of mutations predicted to improve solubility and conformational stability, or one of these properties without affecting the other. The apparent melting temperature of in silico mutants was predicted with NanoMelt ^15^. Of the mutations suggested by this approach, we retained only those that didn’t reduce humanness according to AbNatiV scoring (**Table 7**). These variants were produced in CHO-S cells with production performed at GenScript.

We evaluated these humanized mutants for binding to TfR1 on the cell surface of transiently transfected HEK cells (**Figure 13A**) and found that 13 optimized designs retained binding to cell-membrane TfR1. However, one humanized mutant sample was missing from this experiment—the exhaustive Family D mutant. In particular, all designs that lost binding originated from WT nanobodies that had very unusual residues in the STEM of their CDR3 loop according to the AHo numbering scheme. Such residues were spotted as potential liabilities by the algorithms, and mutations were suggested at their sites. However, we reasoned that the presence of these unusual residues in the WT nanobodies was suggestive of an important role for target engagement, and hence we also made some designs that retained such WT residues (denoted as “WT-CDR3STEM” in **Table 7**). Six out of 7 of these designs retained binding (**Figure 13A**).

**Figure 13.**
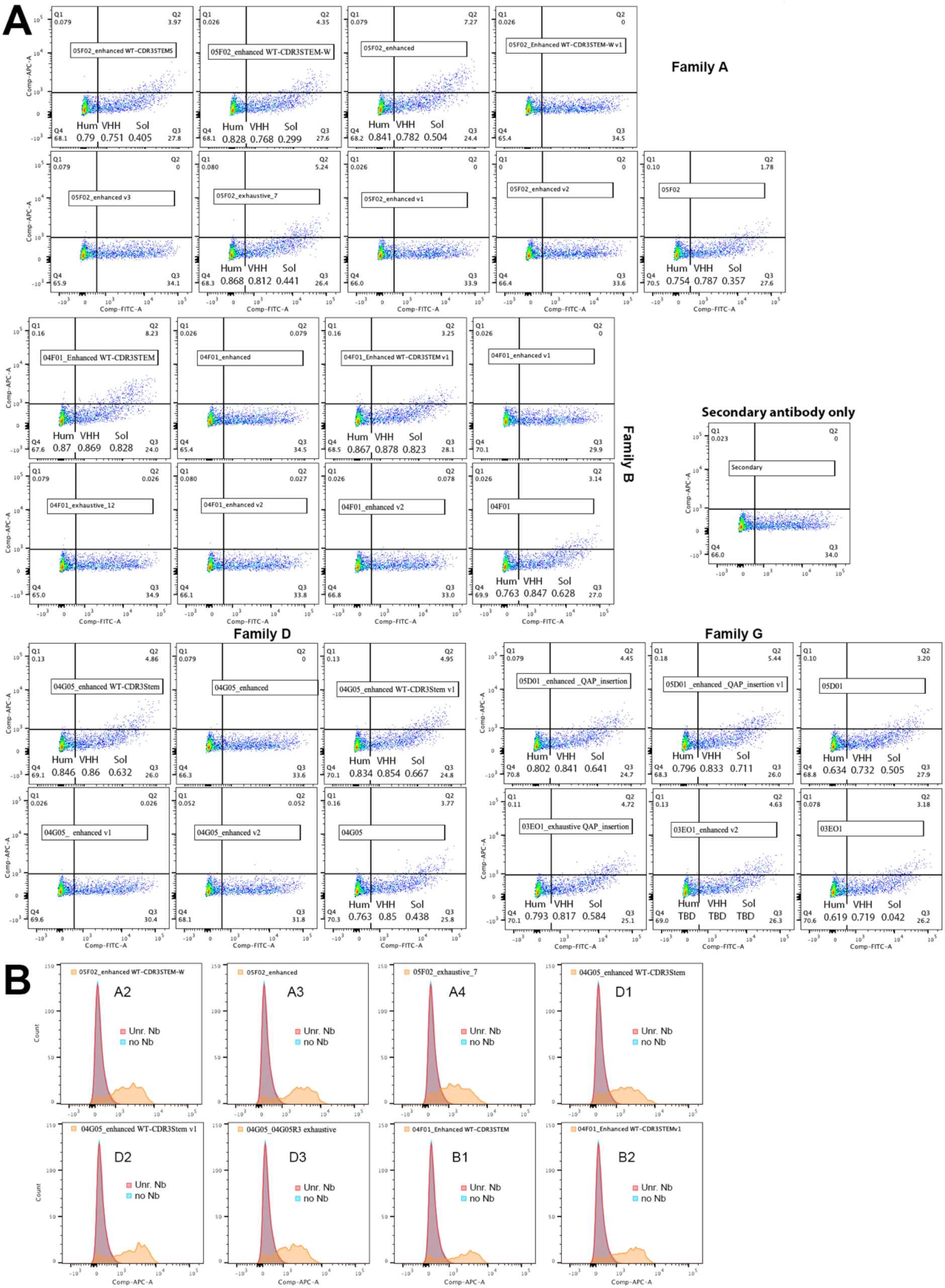
Binding of humanized mutant TfR1b-Nbs to human TfR1. **A)** HEK293 cells were transfected with a vector expressing human TfR1 alongside EGFP. The cells were then treated with each TfR1b-Nb at a concentration of 400 nM, followed by incubation with an anti-His-APC antibody. Secondary antibody staining alone is shown as a negative control. Binding of the wild-type TfR1-Nb from which the mutants originated is also shown for comparison. **B**) HEK293 cells stably expressing human TfR1 were incubated with the eight indicated nanobody mutants (also known as A2, A3, A4, D1, D2, D3, B1 and B2). Two negative controls were included: (1) secondary antibody only (no nanobody), and (2) a non–TfR1-binding nanobody (Unr. Nb). All eight mutants showed robust binding to human TfR1, as compared to the negative controls.

To confirm whether the transiently transfected mutants truly bound human TfR1, we tested seven mutants with optimal drug development properties—including high predicted solubility and humanness (see **Table 7**; mutants A2, A3, A4, B1, B2, D1, and D2)—as well as the Family D exhaustive mutant (renamed D3 in **Table 7**), which had been lost in the previous experiment. All eight mutants were tested on HEK293 cells stably expressing human TfR1. As shown in **Figure 13B**, all bound human TfR1 efficiently.

Consequently, we assessed their BBB permeability in vivo using our *Tfr1* humanized rats. All eight TfR1b-Nbs exhibited high CSF levels, indicating human TfR1-dependent BBB transcytosis (**Figure 14**). Among them, 05F02_enhanced WT-CDR3STEM-W demonstrated the highest CSF/serum ratio. **Table 8** provides details on the age, sex, genotype of the rats used, and the specific nanobodies administered.

**Figure 14.**
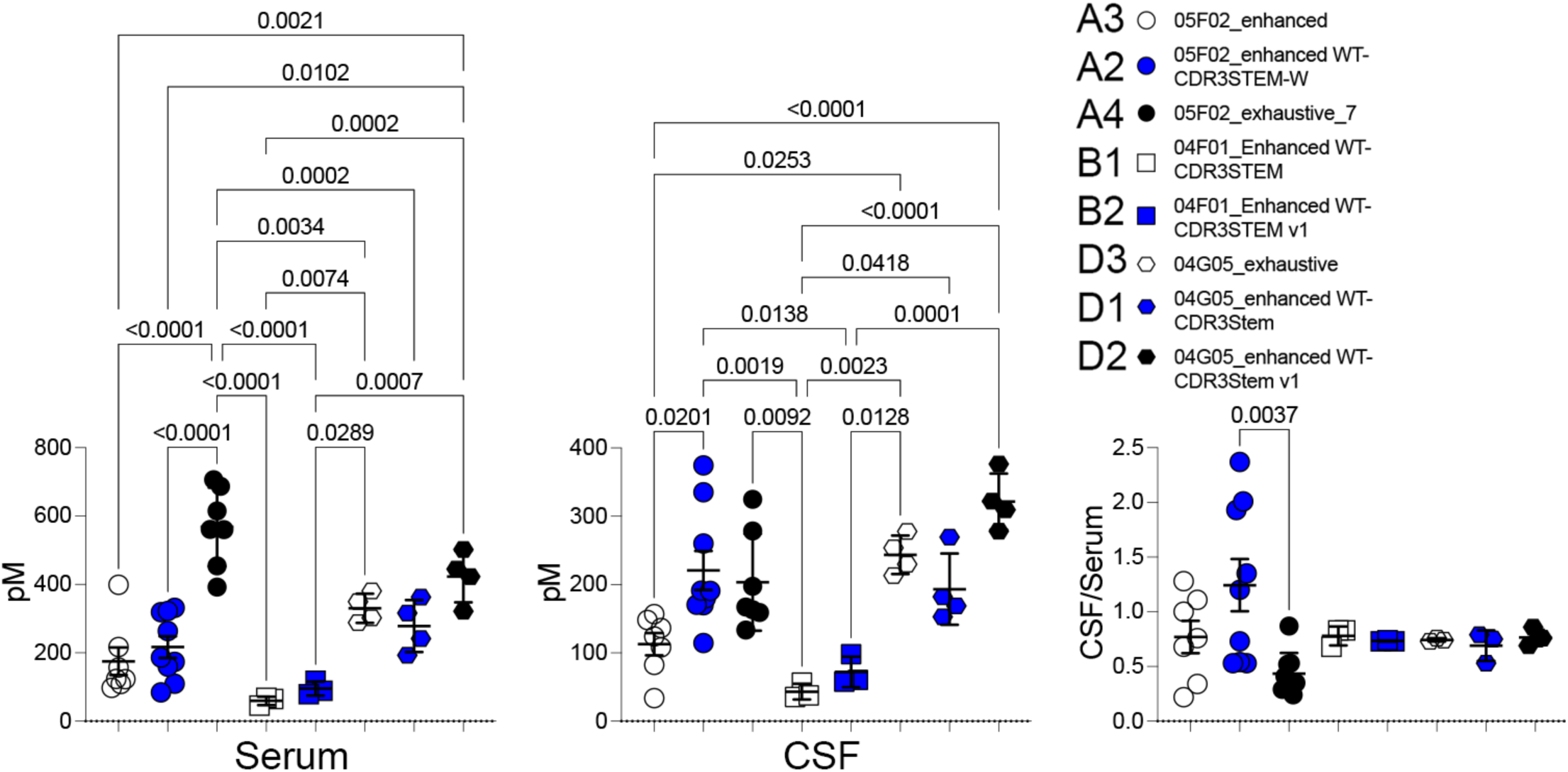
In vivo BBB permeability of eight optimized TfR1b-Nbs in *Tfr1^h/w^*rats. All eight TfR1b-Nbs exhibited elevated CSF levels following IV injection, confirming human TfR1-dependent BBB transcytosis. Among them, 05F02_enhanced WT-CDR3STEM-W showed the highest CSF/serum ratio. Details of the animals and nanobody administration are provided in Table 9.

**Table 8.** Assessment of BBB permeability of optimized TfR1b-Nbs.

| Nanobody | Family | *Tfr1* | sex | Age in days | Time (hrs) | Conc. in Serum [pM] | Conc. in CSF [pM] | CSF/ Ser. |
| --- | --- | --- | --- | --- | --- | --- | --- | --- |
| 05F02 enhanced | A | *h/h* | m | 131 | 16-24 | 122.75 | 156.99 | 1.28 |
| 05F02 enhanced | A | *h/h* | m | 131 | 16-24 | 216.82 | 148.09 | 0.68 |
| 05F02 enhanced | A | *h/h* | m | 100 | 16-24 | 123.96 | 124.15 | 1.00 |
| 05F02 enhanced | A | *h/h* | m | 100 | 16-24 | 97.37 | 108.01 | 1.11 |
| 05F02 enhanced | A | *h/w* | m | 212 | 16-24 | 399.12 | 136.37 | 0.34 |
| 05F02 enhanced | A | *h/w* | f | 212 | 16-24 | 156.72 | 33.77 | 0.22 |
| 05F02 enhanced | A | *h/w* | f | 212 | 16-24 | 108.90 | 82.48 | 0.76 |
| 05F02 enhanced WT-CDR3STEM-W | A | *h/h* | m | 131 | 16-24 | 319.32 | 169.85 | 0.53 |
| 05F02 enhanced WT-CDR3STEM-W | A | *h/h* | m | 131 | 16-24 | 331.44 | 179.84 | 0.54 |
| 05F02 enhanced WT-CDR3STEM-W | A | *h/h* | m | 131 | 16-24 | 322.09 | 170.56 | 0.53 |
| 05F02 enhanced WT-CDR3STEM-W | A | *h/h* | m | 100 | 16-24 | 263.54 | 191.09 | 0.73 |
| 05F02 enhanced WT-CDR3STEM-W | A | *h/w* | m | 82 | 16-24 | 109.81 | 260.22 | 2.37 |
| 05F02 enhanced WT-CDR3STEM-W | A | *h/w* | f | 94 | 16-24 | 173.53 | 335.17 | 1.93 |
| 05F02 enhanced WT-CDR3STEM-W | A | *h/w* | f | 96 | 16-24 | 159.62 | 190.97 | 1.20 |
| 05F02 enhanced WT-CDR3STEM-W | A | *h/w* | f | 94 | 16-24 | 186.12 | 374.53 | 2.01 |
| 05F02 enhanced WT-CDR3STEM-W | A | *h/w* | m | 82 | 16-24 | 84.60 | 114.36 | 1.35 |
| 05F02 exhaustive 7 | A | *h/h* | m | 131 | 16-24 | 615.23 | 324.83 | 0.53 |
| 05F02 exhaustive 7 | A | *h/h* | m | 100 | 16-24 | 560.37 | 197.48 | 0.35 |
| 05F02 exhaustive 7 | A | *h/h* | m | 131 | 16-24 | 687.31 | 278.54 | 0.41 |
| 05F02 exhaustive 7 | A | *h/h* | m | 100 | 16-24 | 392.66 | 133.47 | 0.34 |
| 05F02 exhaustive 7 | A | *h/w* | m | 87 | 16-24 | 561.22 | 163.28 | 0.29 |
| 05F02 exhaustive 7 | A | *h/w* | f | 212 | 16-24 | 454.19 | 159.10 | 0.35 |
| 05F02 exhaustive 7 | A | *h/w* | f | 225 | 16-24 | 706.45 | 167.02 | 0.24 |
| 04F01 Enhanced WT-CDR3STEM | B | *h/w* | m | 92 | 16-24 | 64.86 | 56.40 | 0.87 |
| 04F01 Enhanced WT-CDR3STEM | B | *h/w* | f | 92 | 16-24 | 68.06 | 35.65 | 0.52 |
| 04F01 Enhanced WT-CDR3STEM | B | *h/w* | f | 90 | 16-24 | 45.49 | 37.75 | 0.83 |
| 04F01 Enhanced WT-CDR3STEM v1 | B | *h/w* | m | 92 | 16-24 | 118.29 | 98.00 | 0.83 |
| 04F01 Enhanced WT-CDR3STEM v1 | B | *h/w* | f | 92 | 16-24 | 89.13 | 60.64 | 0.68 |
| 04F01 Enhanced WT-CDR3STEM v1 | B | *h/w* | f | 90 | 16-24 | 79.08 | 57.87 | 0.73 |
| 04G05 exhaustive | D | *h/w* | m | 84 | 16-24 | 381.19 | 277.88 | 0.73 |
| 04G05 exhaustive | D | *h/w* | f | 89 | 16-24 | 288.57 | 213.07 | 0.74 |
| 04G05 exhaustive | D | *h/w* | m | 89 | 16-24 | 302.11 | 229.89 | 0.76 |
| 04G05 exhaustive | D | *h/w* | f | 91 | 16-24 | 349.42 | 253.78 | 0.73 |
| 04G05 enhanced WT-CDR3Stem | D | *h/w* | m | 82 | 16-24 | 362.82 | 269.43 | 0.74 |
| 04G05 enhanced WT-CDR3Stem | D | *h/w* | f | 91 | 16-24 | 316.27 | 168.69 | 0.53 |
| 04G05 enhanced WT-CDR3Stem | D | *h/w* | m | 89 | 16-24 | 241.76 | 182.20 | 0.75 |
| 04G05 enhanced WT-CDR3Stem | D | *h/w* | f | 89 | 16-24 | 192.70 | 152.89 | 0.79 |
| 04G05 enhanced WT-CDR3Stem v1 | D | *h/w* | m | 84 | 16-24 | 321.78 | 278.02 | 0.86 |
| 04G05 enhanced WT-CDR3Stem v1 | D | *h/w* | f | 94 | 16-24 | 445.31 | 309.46 | 0.69 |
| 04G05 enhanced WT-CDR3Stem v1 | D | *h/w* | f | 94 | 16-24 | 423.02 | 322.14 | 0.76 |
| 04G05 enhanced WT-CDR3Stem v1 | D | *h/w* | f | 96 | 16-24 | 502.33 | 376.44 | 0.75 |

**Table 9.**
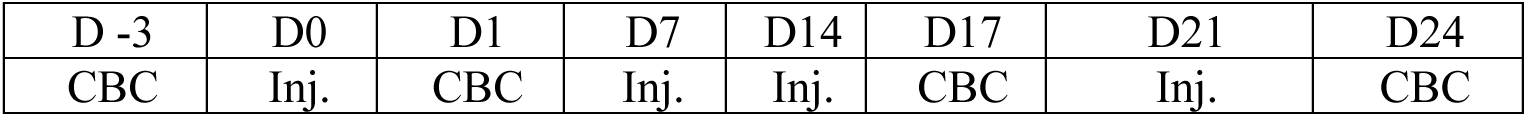
Hematotoxicity assessment. Rats were injected IV on the indicated days (inj.) with either PBS (G1, 3♂/4♀), or TNFI-β-TfR1b-A2 (G2, 3♂/5♀) (1 µL of a 40 µM INN solution in PBS per gram of rat body weight). Complete blood counts (CBC) were performed at several time points: 3 days before the first injection (D-3), 24 hours after the first injection (D1), on Day 17 following three injections, and on Day 24 after four injections. The CBC measurements included white blood cells (WBC), neutrophils (NEU), lymphocytes (LYM), monocytes (MONO), eosinophils (EOS), basophils (BAS), as well as their percentages (NEU %, LYM %, MONO %, EOS %, BAS %), red blood cells (RBC), hemoglobin concentration (HGB), hematocrit (HCT), mean corpuscular volume (MCV), mean corpuscular hemoglobin (MCH), mean corpuscular hemoglobin concentration (MCHC), red cell distribution width (RDW %), platelet count (PLT), and mean platelet volume (MPV).

The eight optimized TfR1b-Nbs were fused to either TNFI-α or TNFI-β—two humanized TNFα inhibitors derived from TNFI-Nb1 ^11^—to generate 16 unique heterodimers (**Figure 15A**). We first evaluated the ability of these heterodimers to bind cell-surface human TfR1, comparing their binding profiles to those of the corresponding parental humanized TfR1b-Nbs used in their construction. As shown in **Figure 15B**, several heterodimers, particularly those incorporating TfR1b-A4, -B1, and -B2, exhibited reduced binding to human TfR1. Whether this reduction can be mitigated by altering the heterodimer design—such as modifying the linker length or sequence—remains to be investigated in future studies.

**Figure 15.**
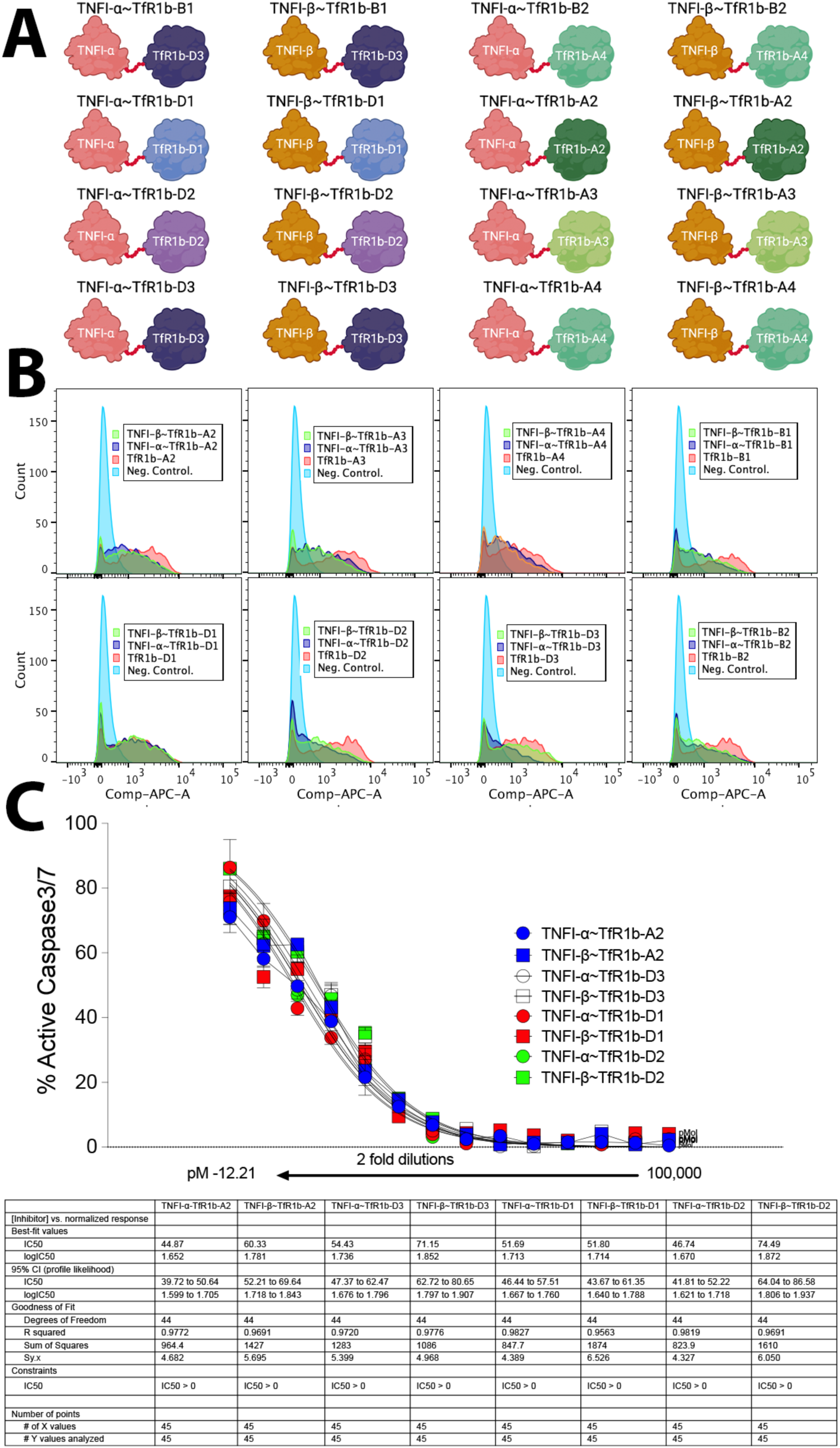
TfR1 binding and TNFα inhibitory activity of optimized and humanized heterodimers. **A**) Schematic representation of the 16 humanized and optimized TNFI–TfR1b heterodimers. **B**) Flow cytometry analysis in HEK293 cells stably expressing human TfR1, comparing the binding of heterodimers to that of their parental humanized TfR1b-Nbs. **C**) Dose-response apoptosis inhibition curves in WEHI-13VAR cells treated with human TNFα and 2-fold serial dilutions of TNFI–TfR1b heterodimers (100,000 to 12.21 pM). Apoptosis was measured using a fluorogenic caspase-3/7 activation assay. The IC₅₀ values of the heterodimers were comparable to those of the parental humanized nanobodies TNFI-α (IC₅₀ = 48.84 pM, range: 42.11–56.56) and TNFI-β (IC₅₀ = 64.31 pM, range: 57.17–72.31).

For the purposes of this study, we prioritized heterodimers containing TfR1b-D1, -D2, -D3, and -A2 for further evaluation. Notably, all selected heterodimers retained TNFα inhibitory activity comparable to TNFI-α and TNFI-β (**Figure 15C**) ^11^—supporting their continued development as therapeutic candidates.

### Evaluating the potential of subcutaneous administration

We selected two heterodimers—TNFI-β-TfR1b-A2 and TNFI-β-TfR1b-D1—for further evaluation of tissue distribution and subcutaneous (SQ) delivery. To test the feasibility of SQ administration—a more patient-friendly alternative to IV, because of its ease of use, steady absorption, reduced clinical resource needs, lower discomfort, and cost efficiency. Approved biologics such as Humira, Enbrel, and Simponi exemplify the success of this route in chronic conditions. In pilot experiments, rats expressing human TfR1 were injected SQ with 1 µL/g of a 40 µM of either TNFI-β-TfR1b-A2 or TNFI-β-TfR1b-D1 solution in PBS. Heterodimers levels were measured 72 hours post-injection. The results are summarized in **Figure 16**. In *Tfr1^h/w^* rats, CSF/serum ratios were 0.14 for TNFI-β-TfR1b-A2 and 0.37 for TNFI-β-TfR1b-D1, with average CSF concentrations of 18.8 pM (TNFI-β-TfR1b-A2) and 5.5 pM (TNFI-β-TfR1b-D1). TNFI-β-TfR1b-A2 achieved higher total systemic levels, while TNFI-β-TfR1b-D1 demonstrated superior BBB permeability. Only the heart (and possibly kidney) showed modest signs of tissue distribution dependent on human TfR1 expression. Importantly, heterodimers were detectable in both serum and CSF three days post-injection, indicating markedly extended in vivo stability compared to conventional nanobodies. Furthermore, the consistently higher serum levels in humanized *Tfr1* rats vs. wild-type rats suggest that target-mediated stabilization contributes to this prolonged half-life.

**Figure 16.**
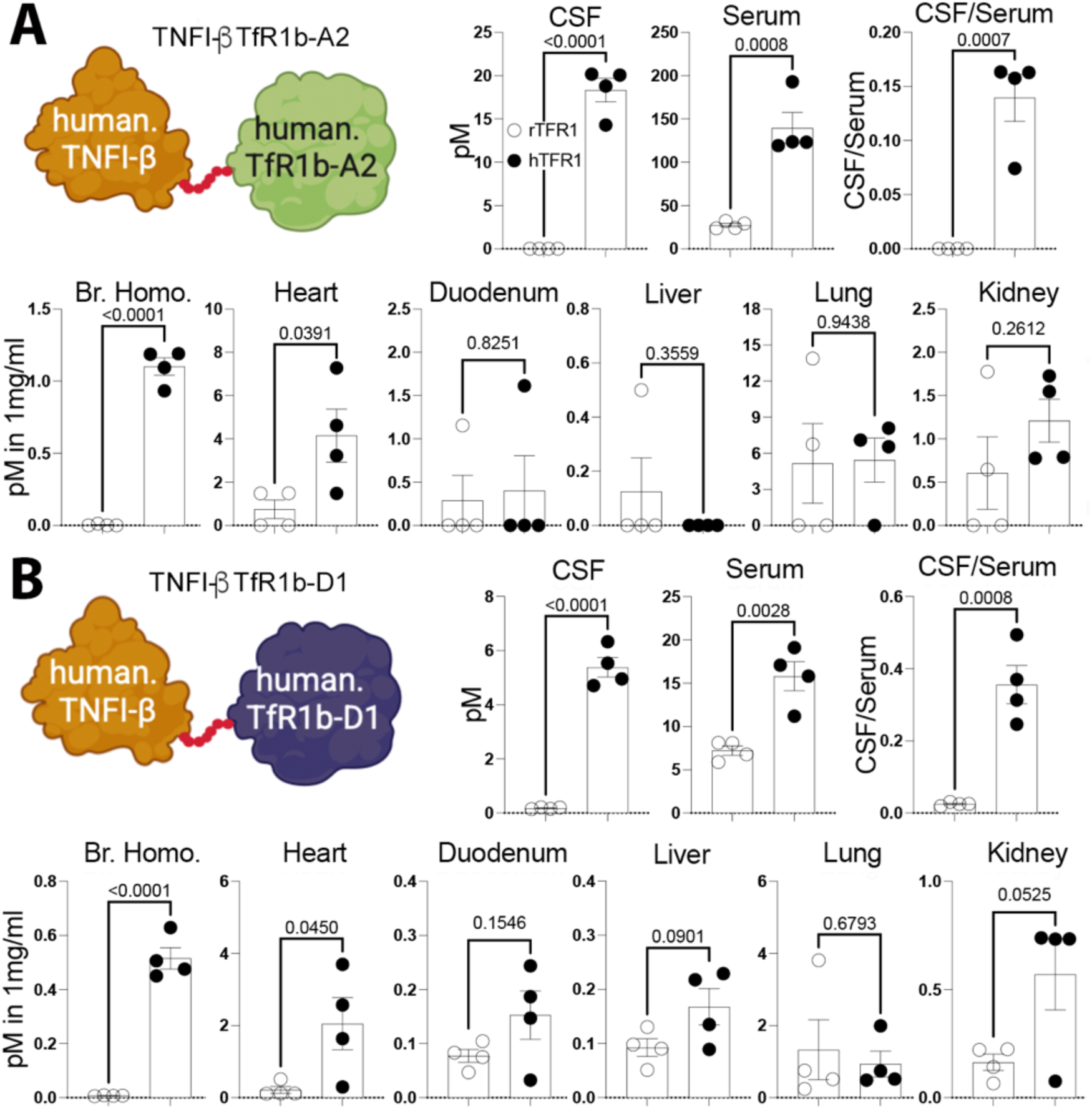
Subcutaneous delivery of humanized heterodimers demonstrates effective BBB transcytosis and extended in vivo stability. *Tfr1^h/w^* and *Tfr1^w/w^* rats were injected subcutaneously with TNFI-β–TfR1b-A2 (**A**) and TNFI-β– TfR1b-D1 (**B**). Heterodimers levels were measured 72 hours post-injection. TNFI-β–TfR1b-A2 achieved higher serum and CSF levels (CSF/serum ratio: 0.14), while TNFI-β–TfR1b-D1 showed superior BBB permeability (CSF/serum ratio: 0.37). Heterodimers remained detectable in both CSF and serum three days post-injection, confirming prolonged in vivo stability. Tissue distribution was primarily brain-specific and human TfR1-dependent, with additional enrichment observed in the heart and, potentially, the kidney. with some enrichment in the heart and, potentially, the kidney.

### Assessing in vivo hematotoxicity

Interfering with TF-TfR1 interaction and/or uptake can have toxic effects, especially anemia. Homozygous null *Tfr1* mice display an embryonic lethal phenotype, while hypo-transferrinemic mice suffer severe anemia ^17^. Although TfR1b-Nabs from Families D and A do not interfere with TF binding or uptake by CHEK-ATP089 cells (**Figure 4** and **5**), we assessed the potential hematotoxicity of heterodimers in vivo. A complete blood count (CBC), conducted by the IRVS core at Rutgers, was performed on ∼4 months-old rats humanized for both TF and TfR1, and administered either TNFI-β-TfR1b-A2 or PBS. This is because hematotoxicity, particularly anemia, is potentially influenced by TfR1 binding. The experimental design is detailed in **Table 9**. The initial CBC (Day-3) established baseline values. Subsequent CBCs (Day 1, Day 17 after three injections, and Day 24 after four injections) monitored for acute and long-term hematotoxic effects. All values remained within normal physiological ranges for rats, indicating that the humanization of *Tf* and *Tfr1* genes has preserved normal blood cell functions. TNFI-β-TfR1b-A2 did not cause significant changes in CBC parameters, including anemia indicators (RBC, HGB, HCT, **Figure 17**), compared to PBS controls. Given that these tests were performed in rats expressing human TF and TFf1, the findings are expected to closely reflect the INNs TNFI-β-TfR1b-A2’ effects in humans, especially regarding holo-TF-TfR1 interactions and iron uptake. This supports the likelihood of low hematotoxicity in humans, particularly given that the therapeutic dosage of heterodimers may be significantly lower than those tested in these experiments.

**Figure 17.**
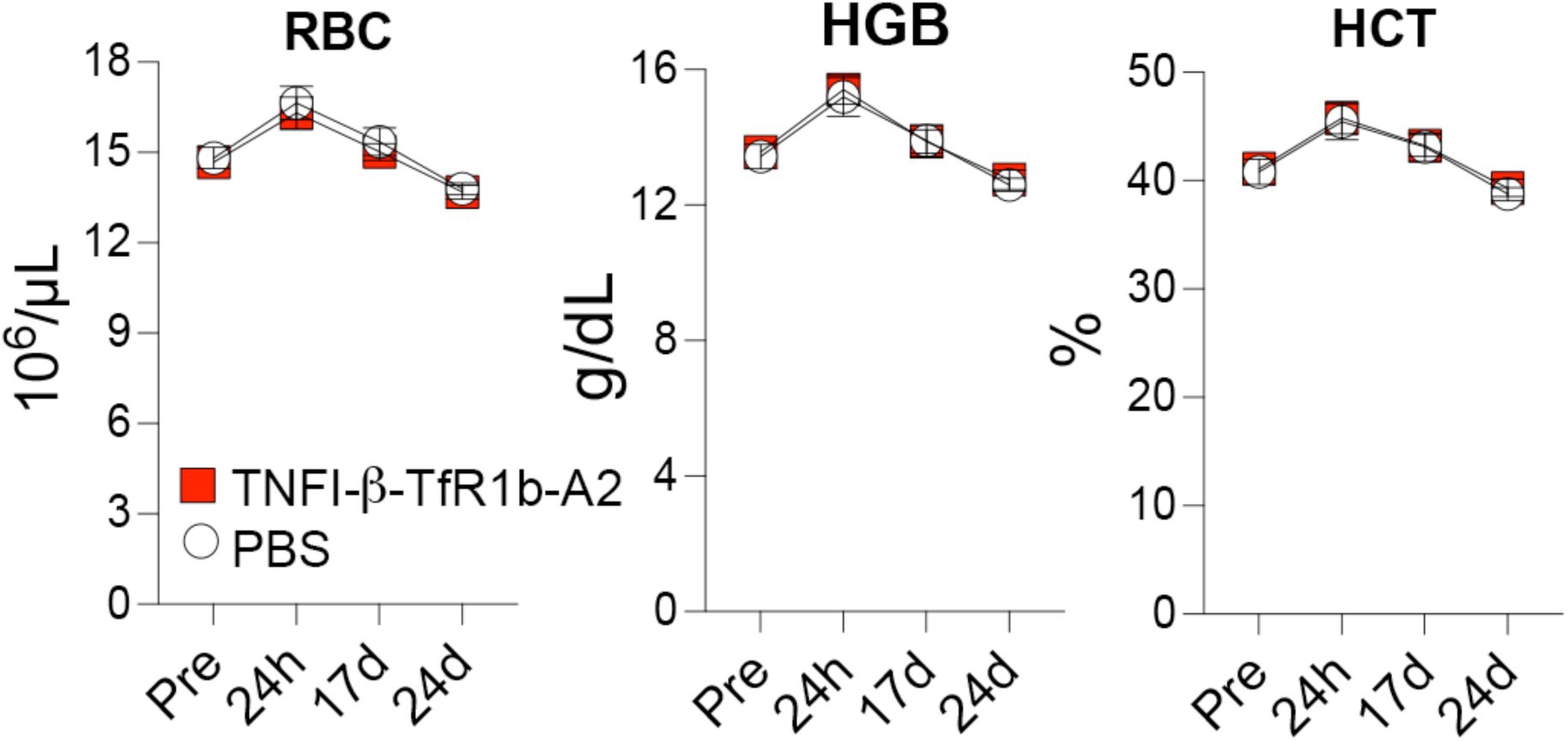
In vivo assessment of hematotoxicity following TNFI-β–TfR1b-A2 administration in humanized rats. CBC analysis on ∼4-month-old rats humanized for both TF and TfR1 (*Tf^h/w^:Tfr1^h/h^*), following administration of TNFI-β–TfR1b-A2 or PBS. No significant changes were observed in key hematological parameters—including RBC, hemoglobin (HGB), and hematocrit (HCT)—compared to PBS-treated controls. All values remained within normal physiological ranges, indicating no detectable hematotoxicity. These results suggest that TNFI-β–TfR1b-A2 is well tolerated in vivo and unlikely to induce anemia through interference with human TF–TfR1–mediated iron transport.

In summary, these results demonstrate that heterodimers retain BBB permeability when administered subcutaneously and are viable for human therapeutic use, combining target specificity, extended stability, and ease of delivery with no detectable adverse effects in *Tf-* and *Tfr1*-humanized rat models.

## Discussion

This study outlines a systematic approach to the generation, selection, optimization, and in vivo validation of humanized TfR1b nanobodies (TfR1b-Nbs) as delivery vehicles for CNS-targeted biologics. The goal was to identify TfR1b-Nbs that bind human TfR1 and exhibit the capacity to cross the BBB in a human TfR1-dependent manner— essential features for their therapeutic translation.

We began with immunization of camelids, cloning and identification of 24 TfR1b-Nbs spanning ten sequence-diverse families. These candidates had been previously validated for their ability to bind human TfR1 on the cell surface. For therapeutic use, production was transitioned from bacteria to CHO-S cells to ensure appropriate folding, post-translational modifications, and reduced immunogenicity—critical parameters for clinical-grade biologics.

Next, we tested each CHO-produced nanobody for its ability to bind cell-surface TfR1, while ensuring that this binding did not interfere with Tf binding or TfR1-mediated iron uptake—key physiological functions of TfR1 that must be preserved for therapeutic safety. Nanobodies that maintained Tf-TfR1 binding and iron transport were then evaluated for their ability to cross the BBB in vivo using newly generated *Tfr1^h^*KI, in which the endogenous rodent TfR1 gene was replaced with the human ortholog. A subset of TfR1b-Nbs demonstrated robust CNS penetration in a human TfR1-dependent manner.

To enhance their clinical potential, the most promising TfR1b-Nbs were subjected to in silico humanization and developability optimization using *AbNatiV* and CamSol Combination. AbNatiV is a machine learning tool that simultaneously evaluates sequence humanness and VHH-nativeness, while CamSol Combination suggests combinations of mutations predicted to improve stability and/or solubility while preserving binding. Mutations were limited to framework regions to preserve antigen-binding activity, and final variants were selected based on high humanness scores, solubility, and retention of structural integrity. Eight optimized TfR1b-Nbs from three families passed this rigorous selection and retained their ability to bind cell-surface TfR1 and cross the BBB in vivo. These were advanced to generate heterodimers with either TNFI-α or TNFI-β—two previously characterized humanized nanobodies targeting TNFα. All 16 resulting heterodimers were initially screened for BBB permeability, and several exhibited favorable pharmacokinetic profiles.

We then focused on two heterodimers—TNFI-β-TfR1b-A2 and TNFI-β-TfR1b-D1—for further characterization, including SQ delivery, a route preferred for chronic therapy due to its convenience, cost-effectiveness, and patient compliance. Remarkably, these heterodimers reached therapeutically relevant concentrations in the CSF and serum 72 hours post-injection, with CSF/serum ratios ranging from 0.14 to 0.37, and significantly extended systemic stability— properties not typical of conventional nanobodies.

Disruption of the TF–TfR1 interaction can result in hematological toxicity, particularly anemia, due to impaired iron uptake. The hematotoxicity study suggest that TfR1b-Nbs are well-tolerated and unlikely to impair iron homeostasis in humans. However, further studies will be required to confirm safety across a broader range of constructs and doses.

These findings validate the humanized TfR1b-Nbs as effective BBB shuttles and demonstrate that their fusion to therapeutic payloads like TNFI-Nbs preserves both BBB permeability and target engagement. Moreover, the ability to deliver these constructs via the SQ route highlights their translational potential for non-invasive, long-term CNS therapy. The pipeline—from immunization to in vivo validation and humanization—is summarized in **Figure 18**.

**Figure 18.**
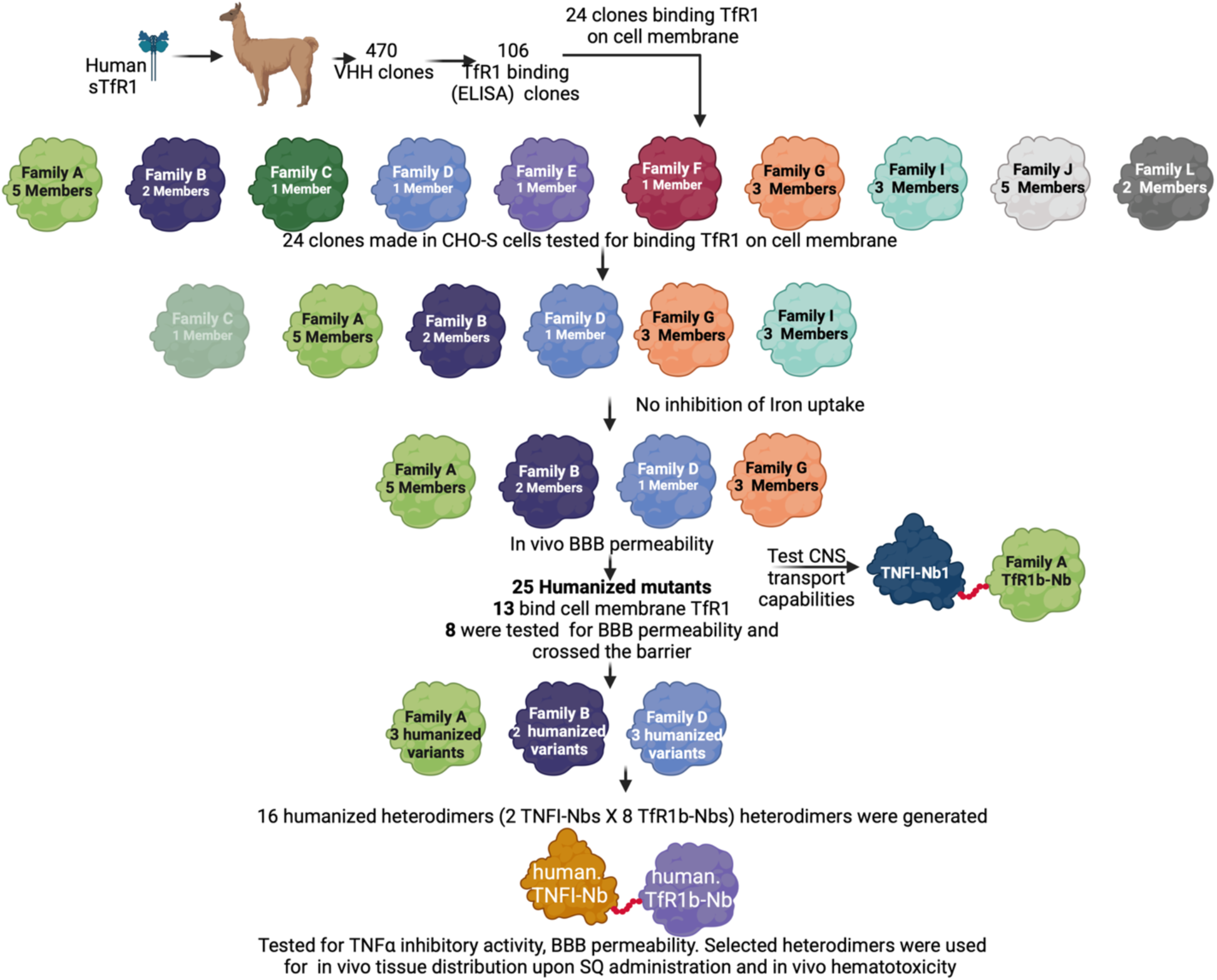
Schematic representation of the workflow for the identification of NewroBus molecules. A step-by-step overview illustrating the process used to discover and optimize NewroBus nanobodies for efficient BBB transcytosis and CNS delivery. The Family C nanobody was not tested.

### Limitations and future directions

While this study establishes humanized TfR1b-Nbs (NewroBus molecules) as highly promising BBB shuttles for therapeutic biologics, several important limitations remain and warrant further investigation.

First, the binding affinities of individual TfR1b-Nbs for human TfR1 have not yet been quantitatively determined. Precise affinity measurements—using techniques such as surface plasmon resonance or biolayer interferometry—will be important to correlate in vitro binding properties with in vivo BBB permeability, serum stability, and CNS accumulation. These data will help refine structure–function relationships and guide further engineering to optimize CNS delivery.

Second, structural studies—such as cryo-electron microscopy (cryo-EM)—would provide valuable insight into how TfR1b-Nbs engage their target at the molecular level. Structural resolution of nanobody–TfR1 complexes could identify precise epitopes, confirm whether the nanobodies bind to one or both subunits of the TfR1 dimer, and whether their binding site is distinct from the transferrin binding site. This information would support the rational design of next-generation molecules that retain TfR1 function while maximizing receptor engagement and transcytosis efficiency.

Third, the cross-reactivity of TfR1b-Nbs with non-human primate (NHP) TfR1 remains to be evaluated. Assessing binding to monkey TfR1 is critical to determine the suitability of NHP models for preclinical safety, pharmacokinetic, and toxicology studies. If cross-reactivity is lacking, alternative strategies may include the development of NHP-compatible surrogate nanobodies or the continued use of rodent models humanized for TfR1 and transferrin.

All tested nanobody constructs, including heterodimers, demonstrated favorable CSF/serum ratios (>0.1), indicative of BBB transcytosis. However, key differences were observed, and a critical question remains: What is the optimal balance between brain selectivity and absolute CNS exposure? Some constructs exhibited high CSF/serum ratios (∼1.0) but relatively lower overall nanobody concentrations in both compartments, while others showed higher absolute levels with more moderate CSF/serum ratios (<0.5). A high CSF/serum ratio may reduce the risk of systemic side effects by limiting peripheral target engagement, whereas high absolute CNS concentrations may be more therapeutically advantageous in diseases requiring potent inhibition of central inflammatory pathways, such as TNFα-driven neuroinflammation.

Furthermore, the relevance of peripheral TNFα inhibition is likely disease-specific. In cases where central inflammation is accompanied by systemic inflammatory components—or where peripheral TNFα contributes to CNS pathology—some degree of peripheral TNFα blockade may be therapeutically beneficial.

In summary, future work will focus on defining nanobody–TfR1 binding affinities, assessing cross-species reactivity, solving nanobody–TfR1 structures, and refining pharmacokinetic and pharmacodynamic profiles. These studies will enable the rational optimization of NewroBus molecules for specific CNS indications and accelerate their progression toward clinical application.

### Conclusion

Taken together, our findings demonstrate that humanized TfR1b-Nbs enable efficient, TfR1-dependent transcytosis across the BBB, delivering otherwise impermeable therapeutic payloads into the CNS. These nanobodies exhibit favorable pharmacokinetics, minimal hematotoxicity, and sustained CNS exposure following subcutaneous administration. Moreover, their ability to localize to astrocytes and microglia highlights their potential for cell-targeted intervention in neuroinflammatory and neurodegenerative diseases. These TfR1b-Nbs have been exclusively licensed to NanoNewron and are collectively referred to as **NewroBus** molecules. As such, NewroBus represents a promising platform for advancing the delivery of biologics to the brain.

## Material sand Methods

Cell lines

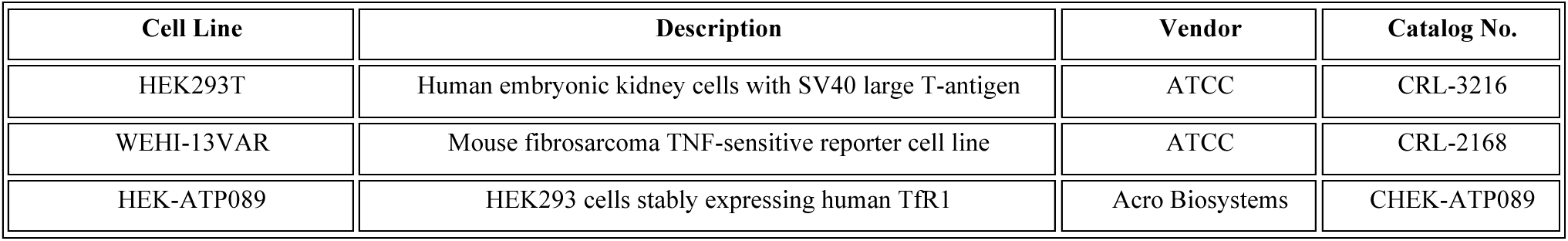

| Cell Line | Description | Vendor | Catalog No. |
| --- | --- | --- | --- |
| HEK293T | Human embryonic kidney cells with SV40 large T-antigen | ATCC | CRL-3216 |
| WEHI-13VAR | Mouse fibrosarcoma TNF-sensitive reporter cell line | ATCC | CRL-2168 |
| HEK-ATP089 | HEK293 cells stably expressing human Tfr1 | Acro Biosystems | CHEK-ATP089 |

Expression constructs

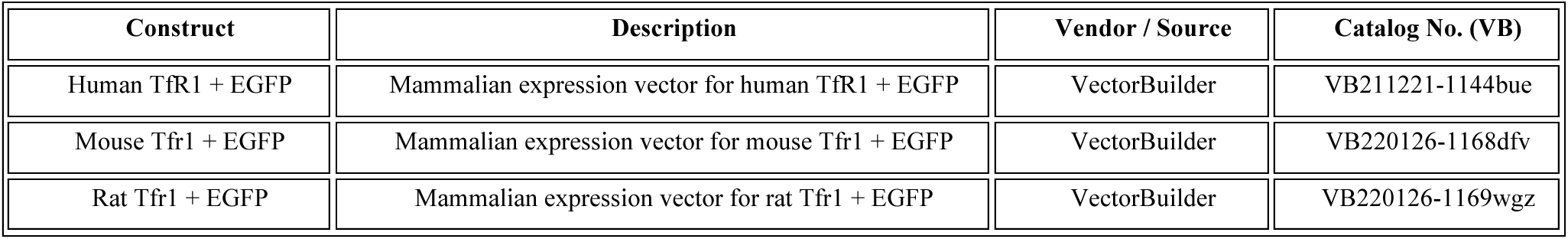

These constructs can be retrieved from VectorBuilder’s plasmid lookup portal: https://en.vectorbuilder.com/design/retrieve.html

Primary Antibodies

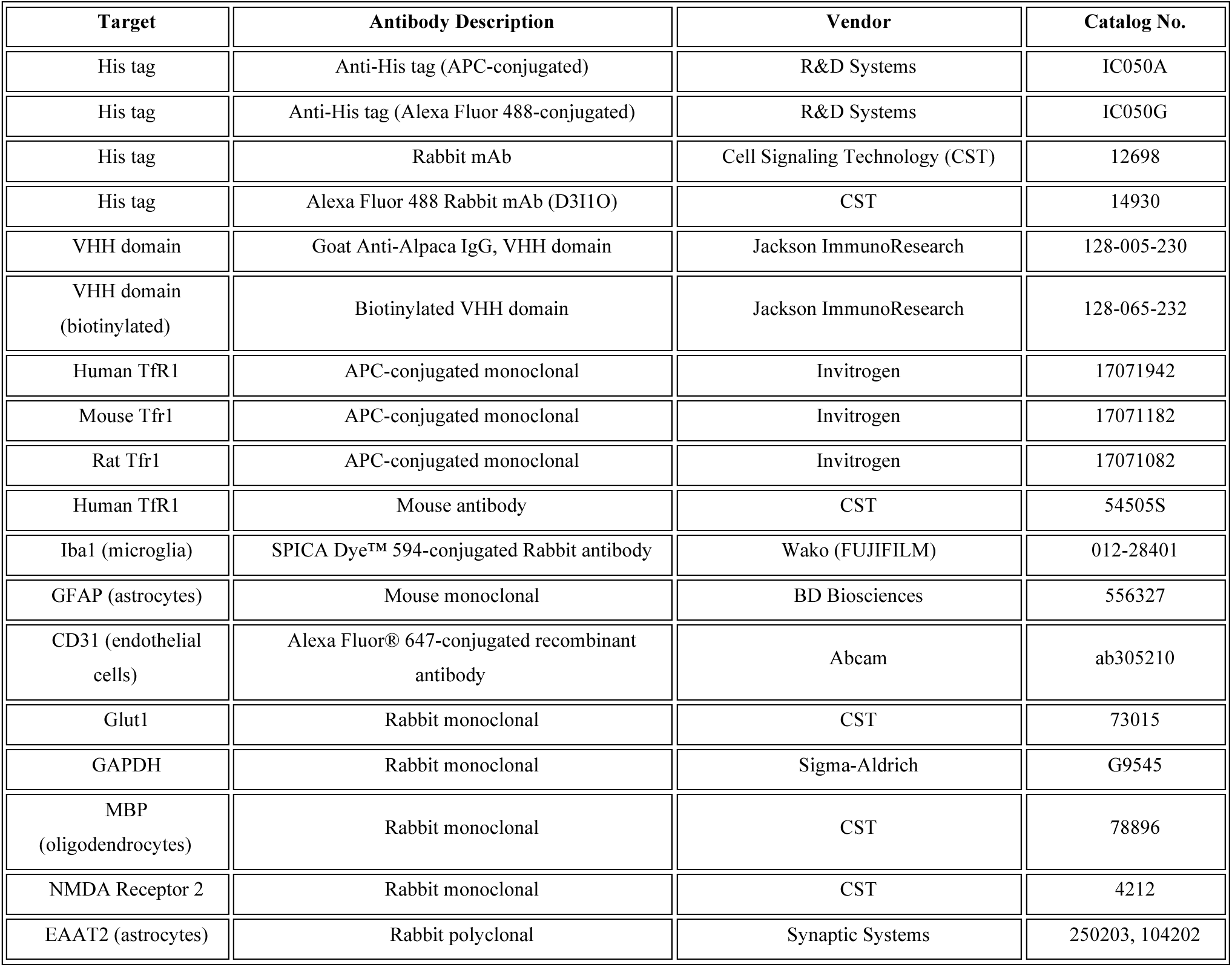

Secondary antibodies

| Target | Antibody Description | Vendor | Catalog No. |
| --- | --- | --- | --- |
| Goat IgG | Anti-Goat IgG Donkey SULFO-TAG | Meso Scale Discovery (MSD) | R32AG |
| Rabbit IgG | Anti-Rabbit IgG Donkey SULFO-TAG | MSD | R32AB |
| Mouse IgG | Goat anti-Mouse IgG (H+L), Alexa Fluor™ 594, HCA | Invitrogen | A-11032 |
| Rabbit IgG | HRP-conjugated secondary antibody | CST | 7074 |
| Rabbit IgG | HRP-conjugated secondary antibody | Southern Biotech | OB405005 |

Recombinant proteins and detection reagents

| Protein / Reagent | Description | Vendor | Catalog No. |
| --- | --- | --- | --- |
| Human Tfr1 | Biotinylated recombinant protein | Acro Biosystems | TFR-H82E5 |
| Human TNFα | Recombinant active trimer | Acro Biosystems | TNA-H5228 |
| Human Transferrin | Unlabeled | Jackson ImmunoResearch | 009000050 |
| Human Transferrin (FITC-conjugated) | Fluorescently labeled | Jackson ImmunoResearch | 009090050 |
| Human Transferrin (pHrodo™ Red-labeled) | pHrodo-conjugated transferrin | Thermo Fisher Scientific | P35376 |
| Biotinylated TNFα | ELISA capture antigen | Acro Biosystems | TNA-H82E3-25ug |
| Biotinylated His Tag | ELISA control | R&D Systems | BAM050 |
| Biotinylated VHH Domain | ELISA control | Jackson ImmunoResearch | 128-065-232 |

General reagents and lab chemicals

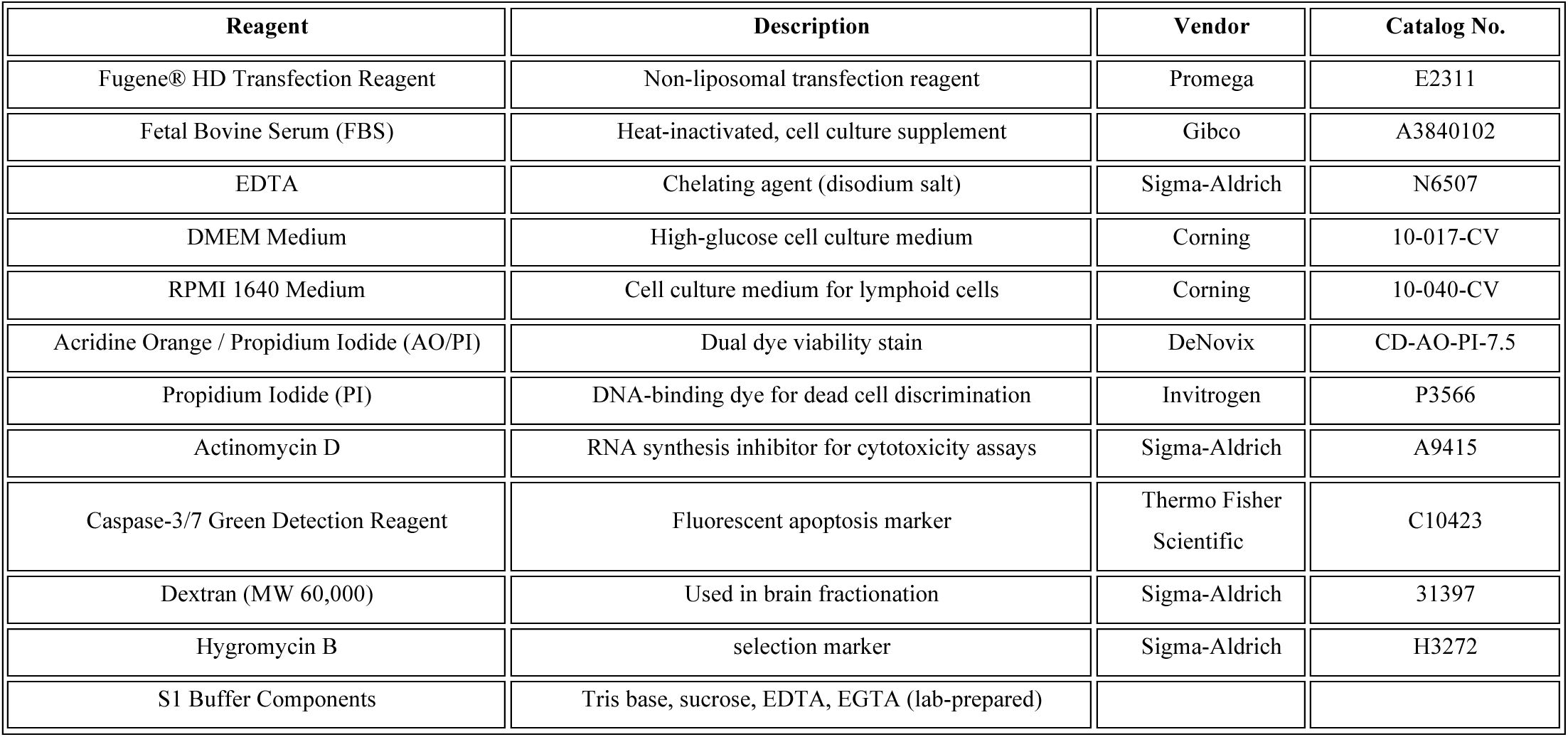

| Reagent | Description | Vendor | Catalog No. |
| --- | --- | --- | --- |
| Fugene® HD Transfection Reagent | Non-liposomal transfection reagent | Promega | E2311 |
| Fetal Bovine Serum (FBS) | Heat-inactivated, cell culture supplement | Gibco | A3840102 |
| EDTA | Chelating agent (disodium salt) | Sigma-Aldrich | N6507 |
| DMEM Medium | High-glucose cell culture medium | Corning | 10-017-CV |
| RPMI 1640 Medium | Cell culture medium for lymphoid cells | Corning | 10-040-CV |
| Acridine Orange / Propidium Iodide (AO/PI) | Dual dye viability stain | DeNovix | CD-AO-PI-7.5 |
| Propidium Iodide (PI) | DNA-binding dye for dead cell discrimination | Invitrogen | P3566 |
| Actinomycin D | RNA synthesis inhibitor for cytotoxicity assays | Sigma-Aldrich | A9415 |
| Caspase-3/7 Green Detection Reagent | Fluorescent apoptosis marker | Thermo Fisher Scientific | C10423 |
| Dextran (MW 60,000) | Used in brain fractionation | Sigma-Aldrich | 31397 |
| Hygromycin B | selection marker | Sigma-Aldrich | H3272 |
| S1 Buffer Components | Tris base, sucrose, EDTA, EGTA (lab-prepared) |  |  |

Consumables and tools for sample collection

| Item | Description | Vendor |
| --- | --- | --- |
| 1 mL Tuberculin Syringe, 25G | For IV injection | Cardinal Health |
| VACUETTE Blood Collection Set, 25G | For tail vein blood collection | Greiner Bio-One |
| BD Microtainer SST Tubes | For tail vein serum isolation | Becton Dickinson |
| BD Vacutainer SST Tubes | For terminal serum isolation | Becton Dickinson |
| Air-Tite Syringe, 5 mL | For cardiac blood collection | Air-Tite Products Co., Inc. |
| Monoject Hypodermic Needle, 21G | For cardiac blood collection | Cardinal Health |
| EXEL Insulin Syringe, 28G | For CSF collection | EXEL |
| Stereotaxic Frame | For CSF collection stabilization | Stoelting Co. |
| MiniCollect K2EDTA Tubes | For CBC analysis | Greiner Bio-One |

Histology and microscopy reagents

| Reagent | Description | Vendor | Catalog No. |
| --- | --- | --- | --- |
| Paraformaldehyde, 4% | Tissue fixative | Electron Microscopy Sciences | 15714-S |
| 30% Sucrose Solution | Cryoprotection | Lab-prepared | — |
| OCT Embedding Compound | Tissue embedding medium | Fisher | 23-730-571 |
| Superfrost Plus Glass Slides | Charged slides for cryosections | Fisher | 22-037-246 |
| Coverslips, 22x50 mm | For mounting stained sections | Corning | 2980-225 |
| Triton X-100 | Detergent for permeabilization | Sigma-Aldrich | T9284 |
| Super PAP Pen | Hydrophobic barrier pen for IHC | IHCWORLD | SPM0928 |
| DAPI Mounting Medium | Antifade mounting with nuclear stain | Southern Biotech | 0100-20 |

Instrumentation and software

| Instrument / Software | Description | Vendor |
| --- | --- | --- |
| MESO QuickPlex SQ 120 | Electrochemiluminescence plate reader | Meso Scale Discovery |
| Incucyte Live-Cell Analysis System | Real-time imaging for cell-based assays | Sartorius |
| Trans-Blot Turbo System | Protein blotting system | Bio-Rad |
| ChemiDoc MP Imaging System | Gel and blot imaging | Bio-Rad |
| Image Lab Software | Western blot quantification | Bio-Rad |
| GraphPad Prism | Data analysis and statistics | GraphPad |
| Biacore 1K SPR System | Surface plasmon resonance (SPR) analysis | Cytiva |
| OpenSPR-XT | Benchtop SPR platform | Nicoya |
| Nikon A1R+ HD Confocal Microscope | Fluorescence imaging with Z-stack acquisition. | Nikon |
| Leica CM1950 Cryostat | Rotary cryostat for the sectioning of frozen tissue specimen | Leica |
| NIS Element Viewer | Confocal image processing | Nikon |

### Generation and screening of camelid anti-TfR1 nanobodies

To generate anti-TfR1 nanobodies (anti-TfR1-Nbs), one alpaca and one llama were immunized with the human TfR1 ectodomain recombinant protein (Acro Biosystems, TFR-H82E5). The immunization protocol began with an initial subcutaneous injection of 0.5 mg TfR1 mixed with Complete Freund’s Adjuvant (CFA) at week 0, followed by booster injections of 0.5 mg TfR1 with Incomplete Freund’s Adjuvant (IFA) every two weeks up to week 14. Serum samples were collected before immunization and at designated time points to assess antibody titers via ELISA, utilizing TfR1-coated plates with appropriate negative and positive controls to ensure specificity.

At weeks 10 and 14, 500 mL of whole blood were drawn from each llama for peripheral blood mononuclear cell (PBMC) isolation. PBMCs were isolated within four hours of blood collection using density gradient centrifugation, ensuring cell viability above 99% as determined by trypan blue exclusion. Total RNA was extracted from the isolated PBMCs using the RNeasy Maxi Kit (Qiagen) and quantified by spectrophotometry, ensuring an A300/A280 ratio greater than 1.9. High-quality RNA was confirmed by agarose gel electrophoresis, displaying distinct 18S and 28S rRNA bands without signs of degradation. Complementary DNA (cDNA) was synthesized from the purified RNA using the SuperScript IV First-Strand Synthesis System (Thermo Fisher Scientific). A nanobody-specific library was constructed by amplifying the variable regions of heavy-chain-only antibodies (VHH) through a two-step PCR process using camelid-specific degenerate primers. The amplified VHH fragments were cloned into the pADL-20c phagemid vector using SfiI restriction sites and transformed into E. coli TG1 cells, resulting in a phage display library containing approximately 2.57 × 10⁹ individual clones. Library diversity was confirmed by sequencing 74 random clones, which revealed 88% contained VHH inserts with intact open reading frames and no duplicate sequences. Library panning was performed against immobilized TfR1 through three rounds of selection to enrich for specific binders. To reduce non-specific interactions, the phage pool was pre-absorbed on BSA-coated wells before each panning round. Enrichment of specific phage binders was monitored using dot assays, which demonstrated increased binding to TfR1 with each successive round.

Following panning, 470 individual clones were screened using an off-phage ELISA to identify those with specific binding to TfR1 and minimal binding to BSA. Positive clones were further validated through repeat screening and sequencing to ensure specificity and diversity. Selected nanobodies were expressed in a non-amber-suppressor strain of E. coli, and periplasmic fractions containing His-tagged nanobodies were purified using His-tag affinity chromatography. Purity was confirmed by SDS-PAGE, and nanobody concentrations were determined by absorbance at 280 nm. Purified nanobodies were dialyzed into PBS (pH 7.4) and filter-sterilized for downstream applications.

### FACS analysis using transfected HEK293T cells and stable HEK293-hTfR1 cells

HEK293T cells (ATCC, CRL-3216) were cultured in DMEM (Corning, 10-017-CV) supplemented with 10% fetal bovine serum (Gibco, A3840102) at 37°C in a humidified 5% CO₂ incubator. For transient expression of transferrin receptor 1 (TfR1), cells were transfected with plasmids encoding human TfR1-EGFP, mouse Tfr1-EGFP, or rat Tfr1-EGFP (VectorBuilder; VB211221-1144bue, VB220126-1168dfv, and VB220126-1169wgz, respectively) using Fugene HD transfection reagent (Promega, E2311), according to the manufacturer’s protocol. Cells were harvested for analysis 24–48 hours post-transfection. HEK293-hTfR1 cells (Acro Biosystems, CHEK-ATP089), a stable clone expressing human TfR1, were maintained under the same conditions and used as a positive control for TfR1 surface expression and nanobody binding. 100μg/mL of Hygromycin B (Sigma, H3272) was used for specific selection of TfR1 expressed clones.

#### Staining procedure

Cells were resuspended in ice-cold FACS buffer (PBS containing 1% BSA and 1 mM EDTA, pH 7.2) and incubated with nanobodies (final concentration 40–100 nM) at 4°C for 45 minutes with gentle shaking. After three washes in FACS buffer, cells were incubated for 30 minutes at 4°C with APC-conjugated anti-His tag antibody (R&D Systems, IC050A) at 1:100 dilution to detect bound His-tagged nanobodies. Following additional washes, cells were stained with propidium iodide (Invitrogen, P3566; 1:1000 dilution) to exclude dead cells.

#### Flow cytometry analysis

Samples were analyzed on a BD LSR Fortessa or similar flow cytometer. Data were gated on live, EGFP-positive cells (for transfection marker) and analyzed for APC signal using FlowJo software. For comparative binding across species-specific TfR1, anti-human, anti-mouse, and anti-rat APC-conjugated antibodies (Invitrogen; Cat. Nos. 17071942, 17071182, 17071082) were also used in parallel at 1:100 dilution.

### TNFα inhibition and IC₅₀ determination using incucyte caspase-3/7 assays

WEHI-13VAR cells (ATCC, CRL-2168), a TNFα-sensitive mouse fibrosarcoma line, were cultured in RPMI 1640 medium (Corning, 10-040-CV) supplemented with 10% fetal bovine serum (Gibco, A3840102) at 37°C with 5% CO₂. Cells were seeded into 96-well plates at 30,000 cells per well in RPMI containing 10% FBS and 1 μg/mL Actinomycin D (Sigma-Aldrich, A9415) to sensitize cells to TNFα-induced apoptosis. Recombinant human TNFα (Acro Biosystems, TNA-H5228) was pre-incubated with varying concentrations of TNFI nanobodies for 30 minutes at room temperature, then added to the cells at a final TNFα concentration of 0.25 ng/mL. Caspase-3/7 activity was monitored using Caspase-3/7 Green Detection Reagent (Thermo Fisher Scientific, C10423) added at a final concentration of 0.5 μM. Plates were immediately placed in the Incucyte Live-Cell Imaging System (Sartorius), and fluorescent signals were monitored every 2–4 hours for up to 24 hours. Fluorescent caspase-3/7 signal intensity was quantified using Incucyte analysis software. TNFα-induced apoptosis was set to 100%, and inhibition was expressed as a percentage of this maximum signal. IC₅₀ values for each TNFI-Nb were calculated using GraphPad Prism with nonlinear regression (log[inhibitor] vs. response – variable slope).

### Transferrin–TfR1 binding competition assay

HEK293-hTfR1 stable cells were cultured in DMEM (Corning, 10-017-CV) supplemented with 10% fetal bovine serum (Gibco, A3840102) and 100μg/mL of Hygromycin B under standard conditions (37°C, 5% CO₂). Cells were harvested at ∼80% confluence, washed with ice-cold PBS, and resuspended in FACS buffer (PBS + 1% BSA + 1 mM EDTA).

#### Competition assay

Cells were incubated for 60 minutes at 4°C in 96-well plates with the following conditions (in 200 µL final volume per well):

### TF-FITC alone

2.5 µM FITC-conjugated human transferrin (Jackson ImmunoResearch, 009090050)

### Nanobody alone

2.5 µM of individual TfR1b-Nbs

### TF-FITC + Nanobody

2.5 µM of both reagents co-incubated.

### TF-FITC + Unlabeled TF

Co-incubation with increasing concentrations of unlabeled human TF (30 nM, 750 nM, 2.5 µM, and 7.5 µM; Jackson ImmunoResearch, 009000050)

Following incubation, cells were washed 3× with FACS buffer and resuspended in buffer containing 1:1000 dilution of propidium iodide (Invitrogen, P3566) to exclude dead cells.

#### Flow cytometry analysis

Fluorescence was measured using a flow cytometer. FITC signal was gated on live cells, and the mean fluorescence intensity (MFI) of the FITC channel was used to quantify TF-FITC binding. Competitive inhibition was determined by comparing MFI between conditions. Decreased MFI in the presence of excess unlabeled TF or TfR1b-Nbs indicated interference with TF-TfR1 binding.

### Transferrin uptake interference assay using incucyte live-cell imaging

HEK293-hTfR1 stable cells were plated in black-walled 96-well tissue culture plates at a density of 30,000 cells per well in DMEM (Corning, 10-017-CV) with 10% fetal bovine serum (Gibco, A3840102) and 100μg/mL of Hygromycin B. After 24 hours, cells reached approximately 80% confluence and were ready for assay. To assess TfR1-mediated uptake, pHrodo™ Red-conjugated human transferrin (Thermo Fisher Scientific, P35376) was used at a final concentration of 312.5 nM. Uptake competition was evaluated by co-incubating with either:

Unlabeled human transferrin at increasing concentrations (30 nM, 150 nM, 750 nM, and 3750 nM), or, selected TfR1b-Nbs at 40 nM.

All reagents were diluted in serum-free DMEM. Cells were incubated at 37°C immediately following reagent addition.

#### Live imaging and quantification

Plates were placed into the Incucyte® Live-Cell Analysis System (Sartorius), and images were acquired every 1–2 hours over a 24-hour period using the red fluorescence channel (excitation/emission ∼560/585 nm). The pHrodo dye emits fluorescence only in acidic intracellular compartments, allowing real-time tracking of transferrin uptake via receptor-mediated endocytosis.

#### Data analysis

Total integrated fluorescence intensity per well was quantified using Incucyte software. Uptake inhibition by TfR1b-Nbs or unlabeled transferrin was calculated as the percent decrease in red signal compared to wells treated with pHrodo-TF alone. All conditions were performed in triplicate and repeated in independent experiments.

### In vivo drug administration, perfusion, and sample collection

#### Intravenous (IV) injection

Rats were briefly warmed under a heat lamp to promote vasodilation. Intravenous drug administration was performed via the lateral tail vein using 1 mL Tuberculin Syringes fitted with 25G needles (Cardinal Health). Nanobodies or fusion constructs were injected slowly at a volume of 1 µL per gram of body weight in sterile PBS to minimize stress and ensure accurate dosing.

#### SQ injection

For SQ administration, animals were gently restrained using a towel, and the compound was injected into the interscapular area using a 1mL Tuberculin syringes fitted with 25G needles (Cardinal Health). Dosing volume was the same as IV: 1 µL of a 40 µM solution per gram of body weight in PBS. Animals were monitored for signs of discomfort or inflammation at the injection site.

#### Perfusion and brain tissue collection

At the designated time points post-injection, animals were deeply anesthetized using isoflurane. For terminal experiments involving brain collection, transcardiac perfusion was performed with cold PBS (without calcium or magnesium) at a flow rate of 10 mL/min for 10 minutes to remove blood from the vasculature. Brains were immediately extracted and processed for either biochemical assays or immunohistochemistry.

### Serum collection

#### Non-terminal

Blood samples (∼300 µL) were collected via tail vein using a 25G VACUETTE Safety Blood Collection Set (Greiner Bio-One) into BD Microtainer Serum Separator Tubes (Becton Dickinson). Samples were allowed to clot for 30 minutes at room temperature, followed by centrifugation at 9000 × g for 30 seconds.

#### Terminal

Blood was collected by cardiac puncture using a 5 mL syringe (Air-Tite Products Co., Inc) fitted with a 21G needle (Cardinal Health), transferred into BD Vacutainer SST tubes, allowed to clot for 30 minutes at room temperature, and centrifuged at 2000 × g for 10 minutes. All serum samples were aliquoted and stored at –80°C.

#### CSF collection

Following transcardiac perfusion and prior to brain tissue collection, CSF was collected via cisterna magna puncture. Animals were placed in a stereotaxic frame (Stoelting Co.). The head was flexed forward, and a midline incision was made at the base of the skull to expose the translucent dura mater over the cisterna magna, using a 28G insulin syringe (EXEL), CSF was slowly withdrawn, carefully avoiding blood contamination. Typically, 50–80 µL of CSF was collected per animal. Samples were snap-frozen in liquid nitrogen and stored at –80°C.

### ELISA for TfR1- and TNFα-based detection of nanobodies. *TfR1-based ELISA*

Streptavidin-coated 96-well plates (Meso Scale Discovery, L15SA or L45SA) were blocked overnight at 4°C with 3% BSA in PBST (PBS + 0.05% Tween-20). Plates were then coated with 0.25 µg/mL biotinylated human transferrin receptor 1 (TfR1) protein (Acro Biosystems, TFR-H82E5) in PBS for 1 hour at room temperature with shaking.

After washing 4× with PBST, samples containing nanobodies were added and incubated overnight at 4°C. Following three PBST washes, wells were incubated with 1 µg/mL Goat Anti-Alpaca IgG, VHH domain (Jackson ImmunoResearch, 128-005-230) for 1 hour at room temperature. After additional washes, SULFO-TAG-labeled anti-goat IgG secondary antibody (Meso Scale Discovery, R32AG; 0.5–1 µg/mL) was added and incubated for 1 hour.

#### TNFα-based ELISA

The same procedure was used with the following modifications: plates were coated with 0.2 µg/mL biotinylated human TNFα (Acro Biosystems, TNA-H82E3-25ug) instead of TfR1. Detection of nanobody binding was performed using either anti-His tag rabbit monoclonal antibody (Cell Signaling Technology, 12698; 1:500 dilution) or Goat Anti-Alpaca IgG VHH domain antibody, depending on the tag configuration. Corresponding SULFO-TAG-labeled anti-rabbit (R32AB) or anti-goat (R32AG) secondary antibodies were used.

#### Plate reading and analysis

After final washes, wells were developed using 2× MSD Read Buffer (Meso Scale Discovery, R92TC) and read on a MESO QuickPlex SQ 120 instrument. Background-subtracted signals were normalized to control wells. TNFI-Nb1 (specific for TNFα) and irrelevant control nanobodies were used to confirm specificity.

### Hematotoxicity Assessment via Complete Blood Count (CBC)

To evaluate hematotoxicity following intravenous administration of TfR1-targeting nanobody constructs, complete blood counts (CBCs) were performed in treated and control rats at multiple time points. Animals received IV injections of either PBS (Group 1, n=7; 3 males, 4 females) or TNFI-β–TfR1b–A2 (Group 2, n=8; 3 males, 5 females) at a dose of 1 µL of a 40 µM solution per gram of body weight. Prior the blood collection, rats were briefly warmed under a heat lamp. Blood samples were collected from the tail lateral vein using a VACUTTE Safety Blood Collection Set (Greiner Bio-One) and transferred into MiniCollect K2EDTA tubes (Greiner Bio-One). Samples were immediately placed on an icepack and kept at +4°C until analysis and transported to the In Vivo Research Services (IRVS) Core Facility at Rutgers University. CBCs were performed using the Heska Element HT5 CBC Analyzer.

CBCs were collected at four time points:

- **Day –3 (D–3):** Baseline, prior to treatment
- **Day 1 (D1):** 24 hours after the first injection
- **Day 17 (D17):** After the third injection
- **Day 24 (D24):** After the fourth injection

The following parameters were measured:

***White blood cell (WBC) profile:***

- Total WBC (×10³/μL)
- Neutrophils (NEU), Lymphocytes (LYM), Monocytes (MONO), Eosinophils (EOS), Basophils (BAS)
- Percent distribution: NEU%, LYM%, MONO%, EOS%, BAS%

***Red blood cell (RBC) profile:***

- RBC count (×10⁶/μL), Hemoglobin (HGB, g/dL), Hematocrit (HCT, %)
- Mean corpuscular volume (MCV, fL), Mean corpuscular hemoglobin (MCH, pg)
- Mean corpuscular hemoglobin concentration (MCHC, g/dL)
- Red cell distribution width (RDW, %)

***Platelet profile:***

- Platelet count (PLT, ×10³/μL), Mean platelet volume (MPV, fL)

These data were used to monitor for treatment-related hematologic changes, particularly indicators of anemia, leukocyte shifts, and thrombocytopenia.

### Fractionation of brain tissue

To evaluate the distribution of nanobodies between the brain vasculature and parenchyma, rats were deeply anesthetized and perfused transcardially with cold PBS. The right hemisphere was dissected, and choroid plexuses were removed. Tissue was homogenized in 5 mL of ice-cold S1 buffer (250 mM sucrose, 20 mM Tris-base pH 7.4, 1 mM EDTA, 1 mM EGTA) using a glass-Teflon homogenizer (10 strokes). Homogenates were centrifuged at 1,000 × g for 10 minutes at 4°C. Pellets were resuspended in 2 mL of 17% dextran (MW 60,000; Sigma, 31397) and centrifuged at 4,200 × g for 15 minutes. The pellet was collected as the capillary-enriched vasculature fraction. The supernatant was diluted in S1 buffer and centrifuged again at 4,200 × g for 15 minutes. The resulting pellet was collected as the vascular-depleted parenchymal fraction.

Both fractions were lysed in ice-cold S1 buffer containing protease and phosphatase inhibitors and sonicated (50% amplitude, 3 × 10 s bursts with 30 s interval rests). Protein concentrations were determined by Bradford assay.

Aliquots were used for ELISA and Western blot.

### Western Blotting of Brain Fractions

Equal amounts of protein from homogenate, vasculature, and parenchymal fractions were mixed with LDS sample buffer (Invitrogen, NP0007) containing 10% β-mercaptoethanol, heated at 95°C for 5 minutes, and loaded onto 4–12% Bis-Tris polyacrylamide gels (Bio-Rad, 3450125). Electrophoresis was followed by transfer to nitrocellulose membranes using the Trans-Blot Turbo System (Bio-Rad) at 25 V for 7 minutes. Membranes were blocked with 5% non-fat milk (Bio-Rad, 1706404) in PBST (PBS + 0.05% Tween-20) for 45 minutes, then incubated overnight at 4°C with primary antibodies (1:1000 dilution in blocking buffer):

- Anti-human TfR1 (CST, 13113)
- Anti-Glut1 (CST, 73015) – endothelial cell marker
- Anti-GAPDH (Sigma, G9545) – loading control
- Anti-VAMP2, NMDAR2B (CST, 4212) – neuronal markers
- Anti-IBA1 (Wako, 01620001) – microglial marker
- Anti-EAAT2 (Synaptic Systems, 250203 or 104202) – astrocytic marker
- Anti-MBP (CST, 78896) – oligodendrocyte marker

After washing, membranes were incubated for 45 minutes at room temperature with HRP-conjugated secondary antibodies (anti-rabbit: CST 7074 or Southern Biotech OB405005, 1:1000 in 5% milk). Detection was performed using Clarity ECL substrate (Bio-Rad, 1705061), and bands were visualized with the ChemiDoc MP Imaging System (Bio-Rad). Densitometry analysis was performed using Image Lab software.

These blots validated the separation of vascular and parenchymal compartments and confirmed human TfR1 expression and nanobody localization across fractions.

### Immunohistochemistry (IHC)

Rats were deeply anesthetized and perfused transcardially with PBS (without calcium and magnesium), followed by fixation with 4% paraformaldehyde (PFA; Electron Microscopy Sciences, 15714-S) in PBS at a flow rate of 10 mL/min for 10 minutes. Brains were extracted and post-fixed in 4% PFA at 4°C for 24 hours on a shaker. After fixation, brains were washed twice in PBS and incubated in 30% sucrose in PBS at 4°C for 48 hours for cryoprotection. Brains were embedded in OCT compound (Fisher, 23-730-571), frozen, and stored at –80°C. Coronal sections (20 μm thick) were cut on a cryostat (Leica CM1950) and mounted onto charged glass slides (Fisher, 22-037-246), then stored at –80°C until use.

#### Immunostaining procedure

Slides were brought to room temperature, and a hydrophobic barrier was drawn around each section using a Super PAP Pen. Sections were rehydrated in PBS for 10 minutes and blocked in 10% normal goat serum containing 0.3% Triton X-100 (Sigma, T9284) for 1 hour at room temperature.

Sections were incubated overnight at 4°C with the following primary antibodies diluted in 5% serum with 0.3% Triton X-100:

| Primary Antibody | Target | Dilution | Vendor / Cat. No. |
| --- | --- | --- | --- |
| His-tag (Alexa Fluor 488) | Nanobody localization | 1:250 | CST, 14930 |
| Anti-IBA1 (SPICA Dye 594) | Microglia | 1:250 | Wako, 012-28401 |
| Anti-GFAP | Astrocytes | 1:250 | BD Biosciences, 556327 |
| Anti-human TfR1 | Human TfR1 | 1:250 | CST, 54505S |
| Anti-CD31 (Alexa Fluor 647) | Endothelial cells | 1:250 | Abcam, ab305210 |

After three washes in PBS containing 0.3% Triton X-100, sections were incubated for 2 hours at room temperature with appropriate fluorescent secondary antibodies diluted 1:1000 in PBS with 5% serum and 0.3% Triton X-100. The following secondary antibody was used:

| Secondary Antibody | Target | Dilution | Vendor / Cat. No. |
| --- | --- | --- | --- |
| Goat anti-Mouse IgG (Alexa Fluor 594) | GFAP, TfR1 | 1:1000 | Invitrogen, A-11032 |

Sections were washed in PBS and mounted using DAPI-containing aqueous mounting medium (Southern Biotech, 0100-20) and cover slipped (Corning, 2980-225).

#### Imaging and analysis

Confocal imaging was performed using a Nikon A1R inverted laser-scanning confocal microscope. Z-stacks and tile scans were acquired using consistent settings across experimental conditions.

Colocalization of nanobody signal with cell-type markers was assessed in the cortex and hippocampus. Image processing was performed using NIS element view.

### Statistical Analysis

All quantitative data are presented as mean ± standard error of the mean (SEM), unless otherwise indicated. Statistical analyses were performed using GraphPad Prism software. Comparisons between two groups were made using unpaired two-tailed Student’s t-tests. For multiple group comparisons, one-way or two-way ANOVA followed by appropriate post hoc tests (e.g., Tukey’s or Sidak’s) were used. A p-value < 0.05 was considered statistically significant. The number of biological replicates (n) is indicated in the figure legends or methods.

