## Supplemental Figures 1-8 for "NewroBus for the brain: humanized TfR1-targeting nanobodies with high BBB permeability and cargo transport capacity" (from another version of this preprint: version 1, 2025-04-24)

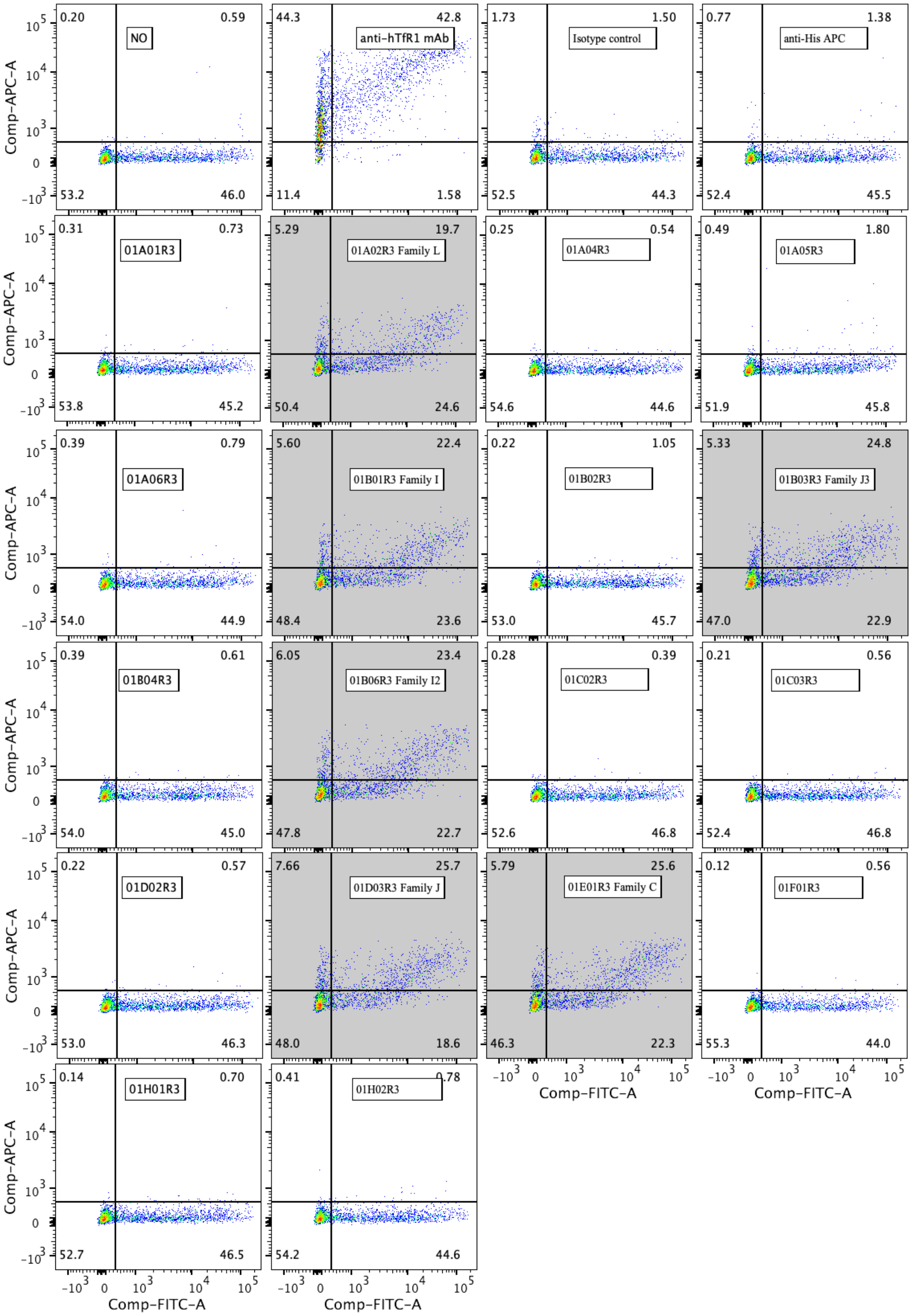


**Figure S1. Screening of 106 bacterially produced anti-TfR1 nanobodies for binding to cell-surface human TfR1**. The first four panels show control staining: No (unstained, negative control); anti-hTfR1–APC antibody (positive control); isotype control–APC (negative control); and anti-His–APC (negative control, secondary antibody used to detect nanobodies). These controls confirm that transfection specifically induces expression of human TfR1 and EGFP. The remaining panels show staining results for the first 22 anti-TfR1 nanobodies tested for binding to human TfR1. The clones binding human TfR1 are in grey.

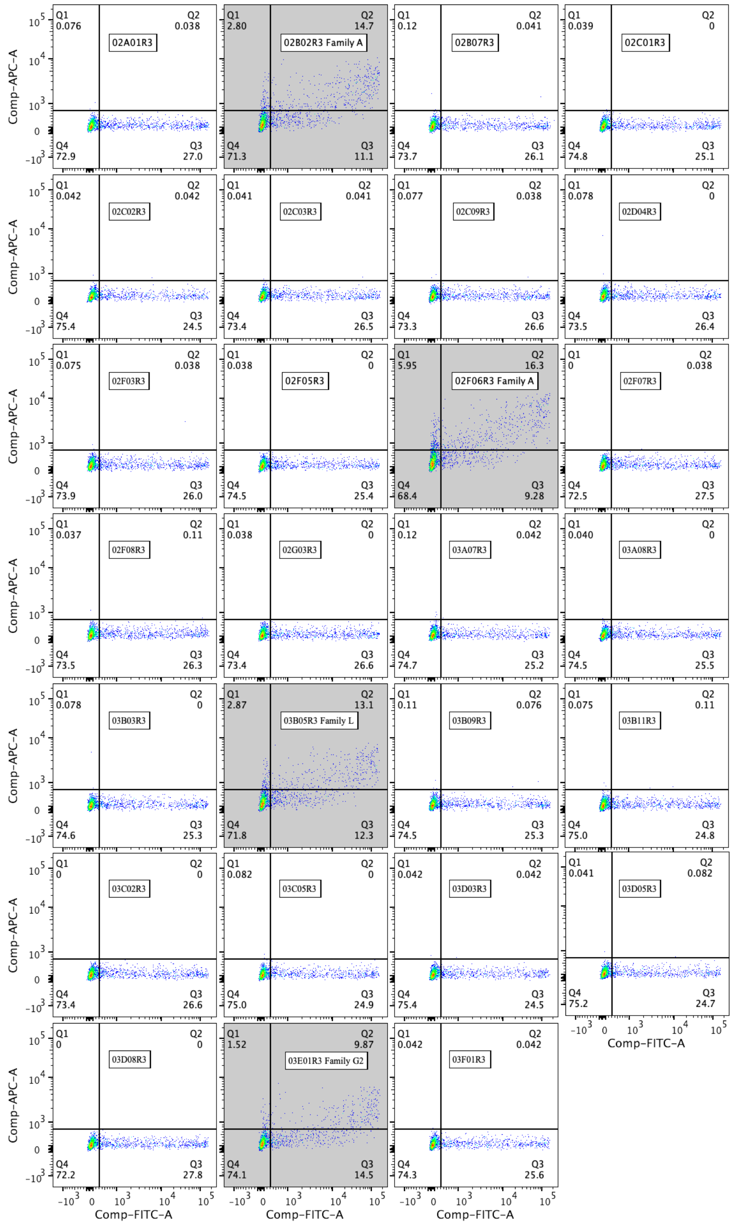

**Figure S2. Screening of 106 bacterially produced anti-TfR1 nanobodies for binding to cell-surface human TfR1**. Staining results of 27 more anti-TfR1 nanobodies tested for binding to human TfR1. The clones binding human TfR1 are in grey.

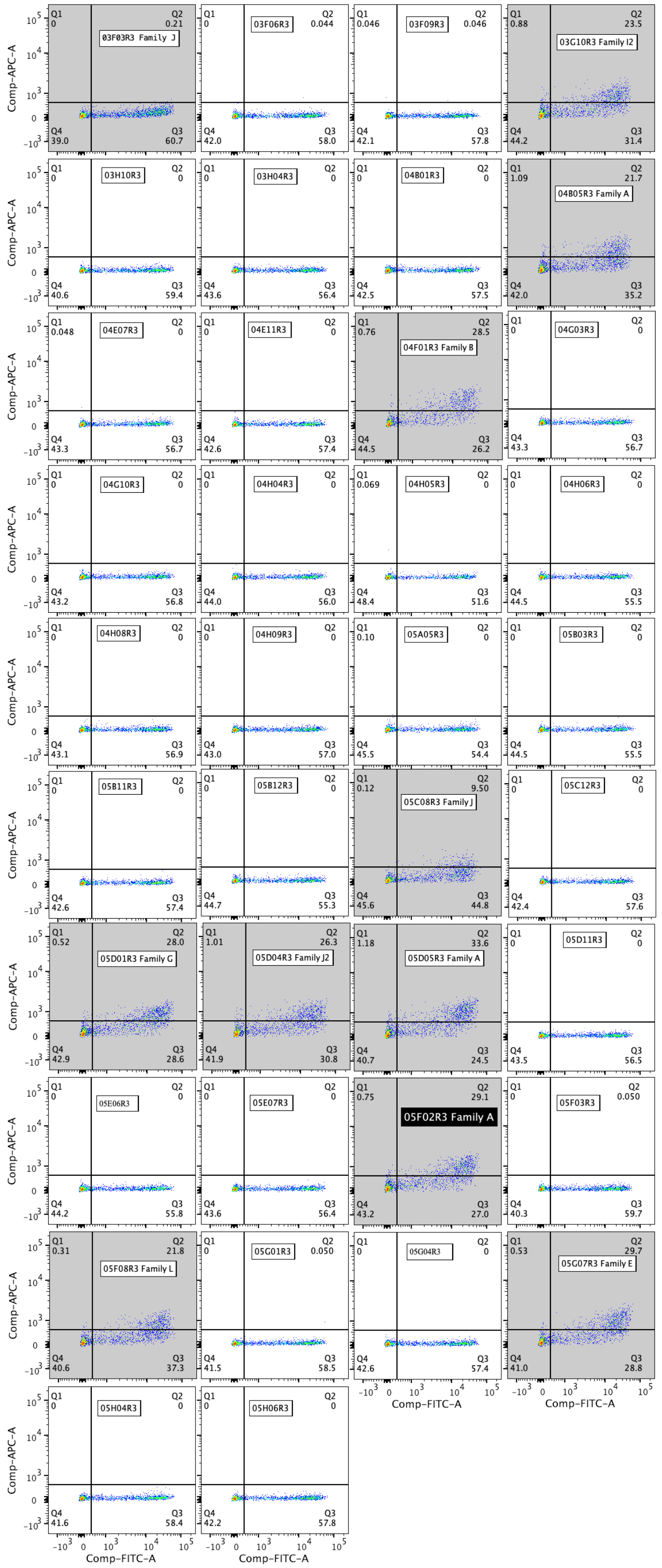


**Figure S3. Screening of 106 bacterially produced anti-TfR1 nanobodies for binding to cell-surface human TfR1**. Staining results of 38 more anti-TfR1 nanobodies tested for binding to human TfR1. The clones binding human TfR1 are in grey. The Nb highlighted in black is one of the two parental sequences from which the final selected NewroBus molecules were derived.

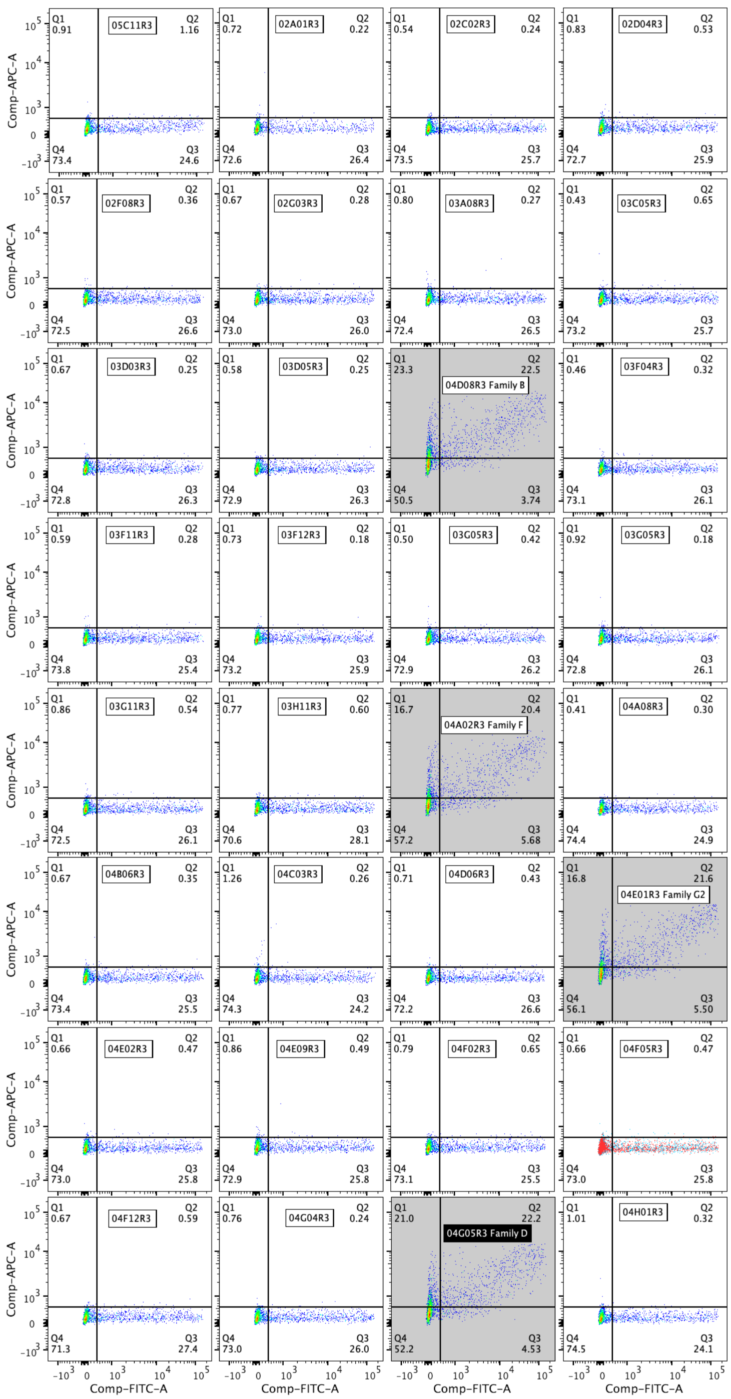

**Figure S4. Screening of 106 bacterially produced anti-TfR1 nanobodies for binding to cell-surface human TfR1**. Staining results of 32 more anti-TfR1 nanobodies tested for binding to human TfR1. The clones binding human TfR1 are in grey. The Nb highlighted in black is one of the two parental sequences from which the final selected NewroBus molecules were derived.

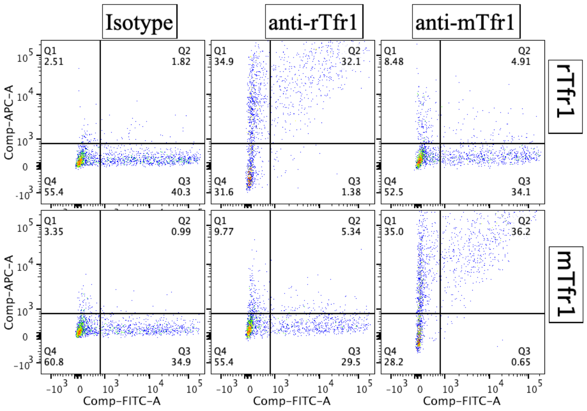


**Figure S5. Transfection of plasmids expressing mouse or rat TfR1 to assess antibody specificity.** HEK293 cells were transfected with plasmids encoding either mouse Tfr1 or rat Tfr1. Specificity of detection was assessed using an isotype control, an anti-mouse Tfr1–APC antibody, and an anti-rat Tfr1–APC antibody. The anti-mouse Tfr1 antibody selectively stained cells transfected with mouse Tfr1, while the anti-rat Tfr1 antibody selectively stained cells expressing rat Tfr1, confirming species-specific binding of each antibody.

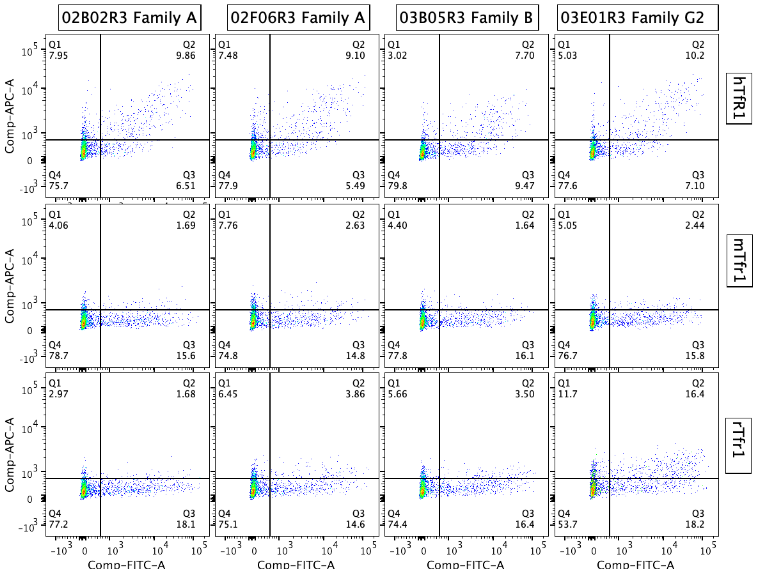


**Figure S6. Part 1: Tfr1b-Nbs recognize human TfR1 but not rat and mouse Tfr1.**

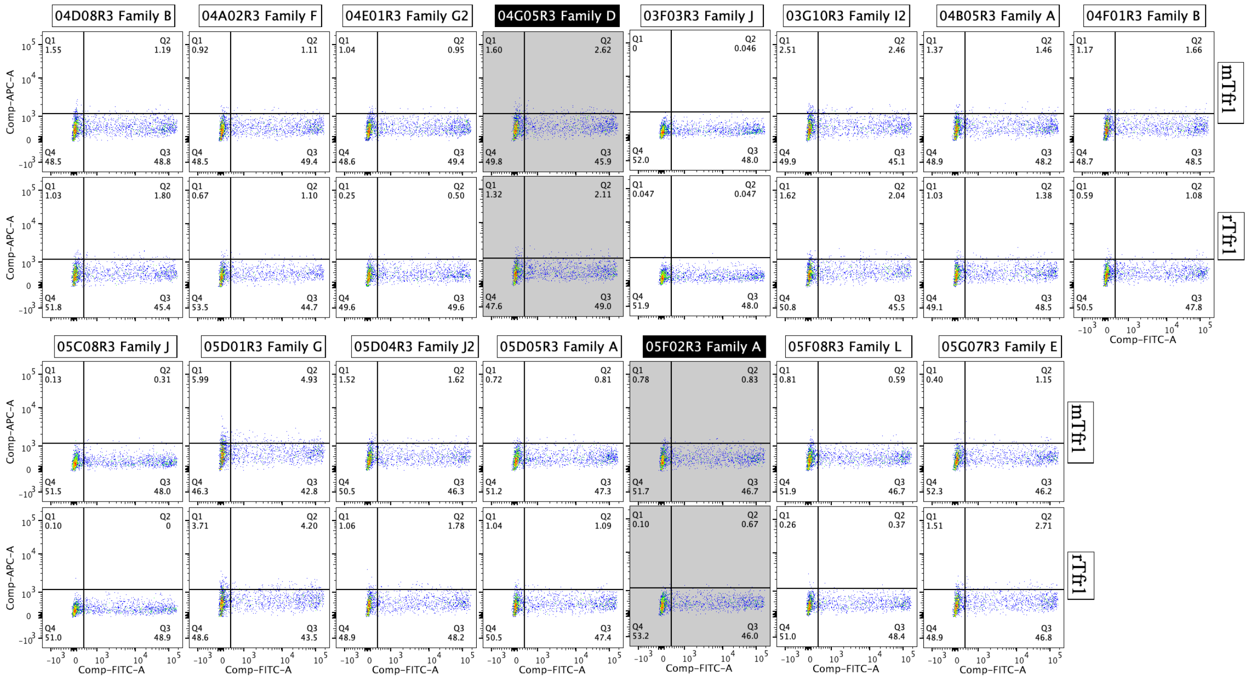


**Figure S7. Part 2: Tfr1b-Nbs do not recognize rat and mouse Tfr1.** The Nbs highlighted in black are the parental sequences from which the final selected NewroBus molecules were derived.

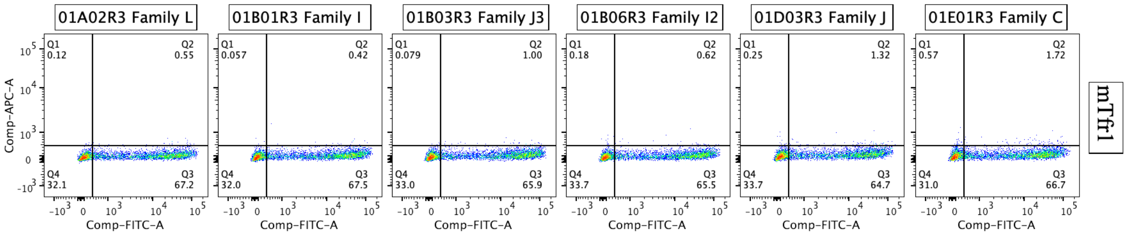


**Figure S8. Part 3: Tfr1b-Nbs do not recognize mouse Tfr1.**
